# Comprehensive multi-omics integration identifies differentially active enhancers during human brain development with clinical relevance

**DOI:** 10.1101/2021.04.05.438382

**Authors:** Soheil Yousefi, Ruizhi Deng, Kristina Lanko, Eva Medico Salsench, Anita Nikoncuk, Herma C. van der Linde, Elena Perenthaler, Tjakko van Ham, Eskeatnaf Mulugeta, Tahsin Stefan Barakat

## Abstract

**Background:** Non-coding regulatory elements (NCREs), such as enhancers, play a crucial role in gene regulation and genetic aberrations in NCREs can lead to human disease, including brain disorders. The human brain is complex and can be affected by numerous disorders; many of these are caused by genetic changes, but a multitude remain currently unexplained. Understanding NCREs acting during brain development has the potential to shed light on previously unrecognised genetic causes of human brain disease. Despite immense community-wide efforts to understand the role of the non-coding genome and NCREs, annotating functional NCREs remains challenging.

**Results:** Here we performed an integrative computational analysis of virtually all currently available epigenome data sets related to human fetal brain. Our in-depth analysis unravels 39,709 differentially active enhancers (DAEs) that show dynamic epigenomic rearrangement during early stages of human brain development, indicating likely biological function. Many of these DAEs are linked to clinically relevant genes, and functional validation of selected DAEs in cell models and zebrafish confirms their role in gene regulation. Compared to enhancers without dynamic epigenomic rearrangement, these regions are subjected to higher sequence constraints in humans, have distinct sequence characteristics and are bound by a distinct transcription factor landscape. DAEs are enriched for GWAS loci for brain related traits and for genetic variation found in individuals with neurodevelopmental disorders, including autism.

**Conclusion:** Our compendium of high-confidence enhancers will assist in deciphering the mechanism behind developmental genetics of the human brain and will be relevant to uncover missing heritability in human genetic brain disorders.

## Introduction

Non-coding regulatory elements (NCREs), such as enhancers, play a pivotal role in gene regulation [1, 2]. Enhancers ensure correct spatio-temporal gene expression, and it is increasingly recognized that genetic aberrations disturbing enhancer function can lead to human disease, including brain disorders [3–6]. Such non-coding genetic variants are expected to explain a considerable fraction of so-called missing heritability (e.g., the absence of a genetic diagnosis despite a high genetic clinical suspicion). These developments are pushing genetic diagnostic investigations to shift from whole-exome sequencing to whole-genome sequencing, and the number of potentially pathogenic non-coding variants found in patients is expected to rise. It is therefore of urgent clinical interest to understand where functionally relevant non-coding sequences are located in the human genome, as this will help to interpret the effects on health and disease.

Despite tremendous progress over the last decades, our understanding of the underlying mechanisms of enhancer biology remains limited due to challenges in annotating functional enhancers genome-wide. Large scale community driven efforts [7–11] and a plethora of individual studies have produced a vast amount of epigenome data sets, such as profiles of histone modifications, chromatin accessibility and chromatin interactions for different human tissues and cell types, that can be used to predict putative enhancers at large scale. More recently, new technologies such as massively parallel reporter assays and CRISPR-Cas9 based screens have entered the stage [12–14], providing novel means to directly test the functionality of non-coding regions. In addition, computational prediction algorithms [15, 16], trained on epigenome and experimental data are improving the capability to predict functional sequences and the effects of variants in these regions.

One of the inherent problems with this increasing amount of data is the difficulty in keeping track of individual data sets and the ability to integrate data from various sources. Usually, individual studies focus on a limited number of cell types or tissues and compare their findings to a small number of previously published data sets. Although this is a logical step, it does not leverage the potential to fine-tune enhancer predictions which integrating all available enhancer data could have. This is illustrated by our previous findings that the overlap between individual enhancer predictions from several studies tends to be quite poor [4]. This is likely caused by heterogeneity of starting biological samples, limitations of current technologies, and differences in data analysis. Although the first two are difficult to change, analysing these data in a similar way could avoid some of the noise and difference generated by data analysis.

Here we undertook such an integrative effort, focusing on human brain development (**Figure 1A, Supplementary Figure 1**). We retrieved virtually all previously published putative enhancers for brain (from Pubmed and enhancer databases, n = ∼1.6 million putative enhancers)([9, 11, 17–29], and performed an integrative analysis of relevant available epigenome data sets (n=494) [8-10, 28, 30-34], after re-analysing the data. Using this approach, we identify around 200 thousand putative critical regions (pCRs) in reported brain enhancers, of which around 40 thousand show dynamic epigenomic rearrangement during fetal brain development, indicating switching on and off of regulatory elements during development. We thus refer to these regions as differentially active enhancers (DAEs). Compared to their non-variable counterparts (nDAEs), DAEs have a higher level of sequence constraint, regulate genes that are expressed during fetal brain development and are associated with brain developmental processes. DAEs are enriched for binding sites of brain relevant transcription factors, brain related GWAS loci and are regulating disease relevant *Online Mendelian Inheritance in Man* (OMIM) genes. We validate a selected number of DAEs using *in vitro* reporter assays and CRISPRi in cell lines, and reporter assays during zebrafish development. Together, this provides an easily accessible and comprehensive resource of NCREs that are likely functional during human brain development.

**Figure 1:**
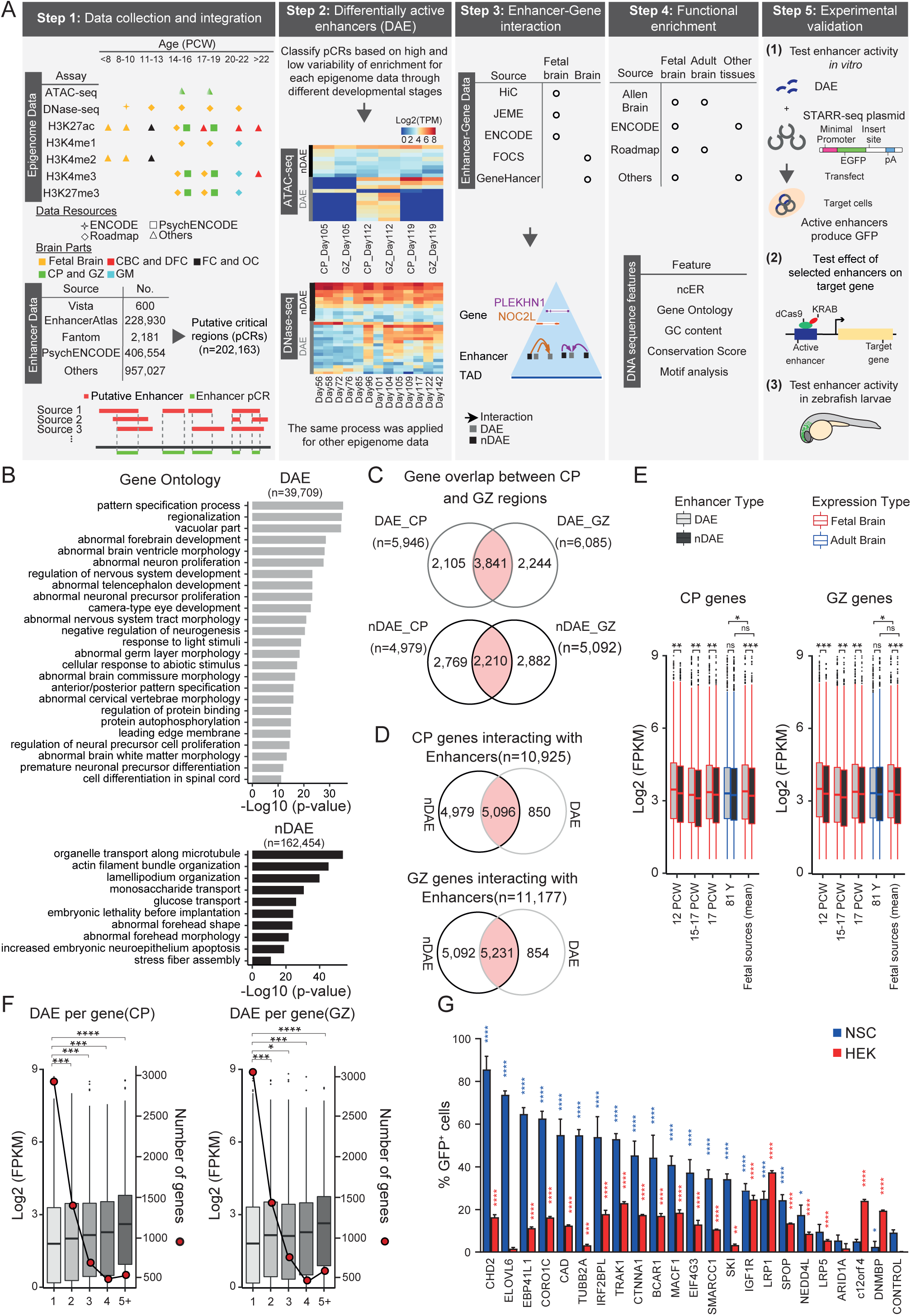
Integrative analysis of brain enhancers during fetal development. A) Various steps taken in the integrative analysis of this study. See text for details. B) Functional enrichment analysis using GREAT [36], for DAEs (upper panel, n=39,708) and nDAEs (lower panel, n=162,454), determined using all pCRs as a background. X-axis reports the −Log10 p-value as determined by GREAT. C) Venn diagram showing the overlap between DAEs (upper panel) and nDAEs (lower panel) interacting with protein coding genes in CP (left) and GZ (right). D) Venn diagram showing the overlap between interactions of protein coding genes with nDAEs (left) and DAEs (right), for protein coding genes in CP (upper panel) and GZ (lower panel). E) Box plots showing gene expression levels as determined by RNA-seq, for genes that interact with DAEs (light gray) or nDAEs (dark gray) in CP (left) and GZ (right), for fetal (red) or adult (blue) brain samples. Boxes are interquartile range (IQR); line is median; and whiskers extend to 1.5 the IQR. PCW, postconceptional week. FPKM, fragments per kilobase of transcript per million mapped reads. * p<0.05; ** p<0.01; *** p<0.001; ns, not significant (wilcox.test). Data obtained from: 12 PCW, Yan et al [105]; 15-17 PCW, De la Torre-Ubieta et al [30]; 17 PCW, Roadmap [10]; 81 years, Roadmap [10]; mean of fetal sources is the mean expression of the first three fetal samples. F) Box plots showing RNA-seq gene expression for genes interacting with 1, 2, 3, 4 or 5 or more DAEs in CP (left) and GZ (right). Left y-axis shows gene expression (log2 FPKM), right y-axis and line plot shows the number of genes per DAE group. * p<0.05; ** p<0.01; ***p<0.001; ****p<0.0001 (wilcox.test). RNA-seq data from Roadmap [10]. G) Bar plot showing the percentage of GFP+ cells in NSCs (blue) and HEK cells (red), from cell transfection experiments with an enhancer reporter plasmid for 22 tested enhancers and an empty plasmid control. Plotted is the percentage of GFP+ in cells co-transfected with an mCherry expressing plasmid, to correct for transfection efficiency. Bars show the average from two independent experiments, with each enhancer tested each in duplicate. Error bars represent standard deviation. * p<0.05; ** p<0.01; *** p<0.001; **** p<0.0001 (one-way ANOVA test followed by multiple comparison test (Fisher’s LSD test).

## Results

### Integrative data analysis identifies differentially active regions during fetal brain development

We started our analysis by collecting relevant fetal brain epigenome data sets and previously published putative enhancers. Epigenome data sets included ChIP-seq for various histone modifications, DNase- and ATAC-seq data from various developmental time points and anatomical regions of human fetal brain, generated by several independent studies, including Roadmap, PsychENCODE and other publications [8-10, 28, 30-34]. All primary data were re-analyzed using identical computational pipelines, and in total we processed 494 data sets. Scrutinizing through previously published literature on enhancers in brain and neuronal cell types, we collected 1,595,292 putative brain enhancers. These included enhancers retrieved from various enhancer databases, such as VISTA, FANTOM and EnhancerAtlas, enhancer predictions from the PsychENCODE consortium, human accelerated regions, ultra-conserved regions and others [9, 11, 17–29]. We first analyzed the overlap between the different putative enhancers, and found only a small overlap between enhancer predictions from different studies. We reasoned that if different enhancer prediction methods used in the individual studies identified the same enhancers that only differ by the exact location or length, by merging the overlaps between different studies we could identify functional relevant parts of enhancers. We thus proceeded to determine putative critical regions (pCRs), by determining the unifying overlaps between all putative enhancers (Step 1 in **Figure 1A**). In this analysis, we kept those putative enhancers that were only found in a single study, merged the overlaps between multiple studies and eliminated those regions that were located within 2 kb upstream and 1 kb downstream of a transcriptional start site (TSS) or which had < 10 reads in epigenome data (see methods). This resulted in 202,163 pCRs, with a total length of 93 Mb, an average size of 460 bps and most pCRs located between 5 and 50 kb away from the closest gene TSS (**Supplementary Figure 2A, B**).

We assumed that enhancers that have functional relevant roles during brain development would show dynamic changes in the levels of histone modifications and chromatin accessibility correlating with their function. To investigate this, we next intersected all pCRs with all epigenome data sets from different time points of fetal brain development and calculated the read count for each pCR region. After TMM-normalization, we performed differential accessibility analysis (for ATAC-seq and DNase data) and generated differential histone modification profiles (for H3K27ac, H3K27me, H3K4me1, H3K4me2, H3K4me3) using edgeR [35]. This resulted in 39,709 pCRs that showed a high variability for these features across developmental timepoints (**Figure 1B, Supplementary Figure 2C, see methods**) which we refer to as differentially active enhancers (DAEs). In contrast, the remaining 162,454 pCRs showed a more constant epigenome pattern and we thus refer to them as not-differentially active enhancers, nDAEs (**Figure 1B, Supplementary Figure 2C**).

Gene ontology analysis using GREAT [36] showed that DAEs were significantly enriched for terms related to brain development, including processes such as pattern specification, neuron proliferation and nervous system development (**Figure 1B**). nDAEs appeared to be enriched for more general terms, including organelle transport and metabolic processes (**Figure 1B**).

To have a more specific view about the genes regulated by these pCRs, we next linked DAEs and nDAEs to their target genes, using different resources, which either link enhancer to gene promoters by direct chromatin interaction as determined by the chromatin conformation capture technique HiC [26] or by predicting enhancer-gene interactions using statistical models and correlation between gene expression, omics data and epigenome features (JEME, FOCS, GeneHancer, ENCODE) [37–40]. Since only a limited number of interactions between DAEs or nDAEs and target genes were supported by >2 of the available resources (**Supplementary Figure 3A**), and as most interactions were predicted by the HiC data, we focused on these HiC predicted ones for the remainder of the analysis. These HiC data were generated from post conceptional week (PCW) 17-18 human brains [26], and were available for the germinal zone (GZ) (containing primarily mitotically active neural progenitors), and the cortical and subcortical plate (CP) (consisting primarily of post-mitotic and migrating neurons).

Taking only those enhancer-promoter interactions that occurred in the same topological associated domain (TAD) into account, we found that from all DAEs, 6,858 and 6,883 for CP and GZ, respectively, interacted with promoters of protein coding genes or lincRNAs, of which the majority of interactions occur with protein coding genes. Similarly, 27,004 and 27,161 nDAEs interacted with target genes in CP and GZ, respectively, with a similar distribution between protein-coding and lincRNAs (**Supplementary Figure 3B, C**).

In total, DAEs interacted with 5,946 and 6,085 protein coding genes in CP and GZ, respectively, of which 3,841 genes were shared between both CP and GZ (**Figure 1C**). The majority of these genes (86%) also had interactions with nDAEs (**Figure 1D**). We next integrated available gene expression data from fetal and adult brain, and found that genes that interacted with a DAE had a significantly higher gene expression compared to those genes not interacting with a DAE, at various regions and stages of fetal development but not in adult brain (**Figure 1E, Supplementary Figure 3D**). In line with earlier findings [41], we find that the more enhancers a gene is interacting with, the higher the gene expression is, and this was also true for the DAEs (**Figure 1F**).

Finally, to validate that DAEs can function as enhancers, we selected 22 DAEs linked to genes, cloned them in an enhancer reporter plasmid [42] and tested their enhancer activity in cell transfection experiments. Upon transfection in human neural stem cells (NSCs), 18 out of 22 tested sequences showed significantly increased percentage of GFP+ cells compared to control (normalized for transfection efficiency using an mCherry spiked-in control), confirming enhancer activity (**Figure 1G**). Transfecting the same plasmids in non-neural HEK cells showed less pronounced activity. This indicates that 81.8% of the tested DAEs had a measurable enhancer activity using this assay in an *in vitro* neural cell type. Of note, this does not exclude activity of those 4 DAEs that do not show enhancer activity in NSCs, in other cell types during fetal brain development.

We conclude that an integrative data analysis of virtually all previously reported brain enhancers identifies a set of DAEs which are associated with a brain developmental gene ontology, increased gene expression in fetal brain and display enhancer activity *in vitro*.

### Multi-gene interacting enhancers regulate genes implicated in multiple cellular processes and have distinguishing sequence characteristics

In order to understand the biological function of DAEs and nDAEs in more detail, we further characterized these two groups. When determining the number of genes that each DAE is interacting with, we found that the majority of DAEs interact with 1 or 2 genes; but, in addition, a considerable fraction of DAEs also interact with more than 2 genes (19.7 % for CP, 19.4% for GZ) (**Figure 2A**), and the same was found for nDAEs (**Figure 2B**). When comparing the enrichment of biological processes for the genes that interact by HiC with these multi-gene interacting DAEs, we found that these genes were enriched for more broad developmental and metabolic processes. However, genes that interact with DAEs that only regulate single genes were enriched for more specific brain related terms, such as “neuron differentiation” and “neuron migration”. Similar results were obtained using GREAT analysis, where multi-gene interacting DAEs for example were enriched in mouse phenotypes associated with “early lethality”, whereas DAEs associated with only a single gene were enriched for “regulation of neural precursor cell proliferation”.

**Figure 2:**
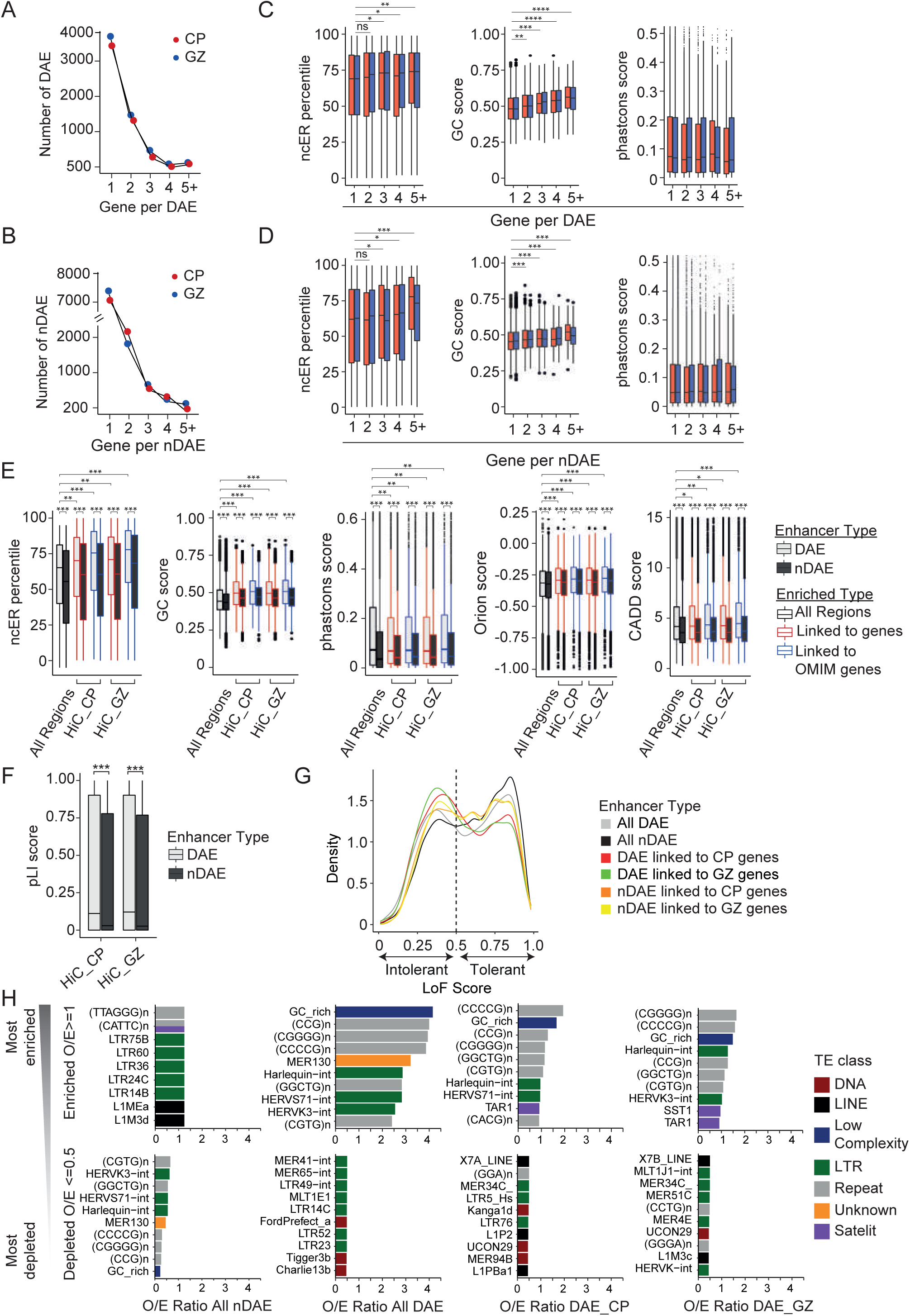
Distinct sequence characteristics between DAEs and nDAEs. A) Line graph showing the number of protein coding genes (1, 2, 3, 4 or 5 or more) that each DAE is interacting with, and the number of DAEs per categorie, for CP (red) and GZ (blue). B) As A), but here for nDAEs C) Box plots showing the median ncER percentile (left) [43], GC content score (middle) [44] or phastcons score (right) [44] for DAEs-CP (red) and -GZ (blue) that interact with 1, 2, 3, 4 or 5 or more protein coding genes. Boxes are IQR; line is median; and whiskers extend to 1.5 the IQR. * p<0.05; ** p<0.01; *** p<0.001; **** p<0.0001; ns, not significant (wilcox.test). D) As C), but here for nDAEs E) Box plots, showing from left to right ncER percentile [43], GC content score [44], phastcons score [44], Orion score [47] and CADD score [48], for all DAEs (light gray) and nDAEs (dark grey), or for those DAEs and nDAEs that are interacting in CP or GZ with protein coding genes (red) or genes with a known OMIM phenotype (blue). Boxes are IQR; line is median; and whiskers extend to 1.5 the IQR. * p<0.05; ** p<0.01; *** p<0.001; ns, (wilcox.test). F) Box plot showing the pLI score [49] of genes interacting with DAEs (light gray) and nDAEs (dark gray) in CP or GZ. Boxes are IQR; line is median; and whiskers extend to 1.5 the IQR. * p<0.05; ** p<0.01; *** p<0.001; ns, (wilcox.test). G) Kernel density plot showing the distribution of loss-of-function tolerance scores for non-coding sequences [50] for all DAEs (light gray), all nDAEs (dark gray), DAEs linked to protein coding genes in CP (red), DAEs linked to protein coding genes in GZ (green), nDAEs linked to protein coding genes in CP (orange), and nDAEs linked to protein coding genes in GZ (yellow). H) Bar chart showing the 10 most enriched (upper panel) and the 10 most depleted (lower panel) transposable elements (TEs) overlapping with from left to right all nDAEs, all DAEs, DAEs interacting with protein coding genes in CP and DAEs interacting with protein coding genes in GZ. Plotted is a ratio between the observed (O) number of TEs over the expected (E). Different classes of TE are indicated with different colors as indicated.

We next asked whether DAEs that regulate single or multiple genes could have distinguishing DNA sequence characteristics, which could support their presumed distinct functional roles. To answer this, we focused on scores that provide some weight based on the underlying sequences: non-coding essential regulation (ncER) score [43], GC content [44] and phastcons score [44]. The ncER scores were recently established using a machine learning model [43], taking functional, mutational and structural features into account, including sequence constraint in the human population, and provides a score where 0 is non-essential, and 1 is putative-essential. We observe that DAEs that interact with 3 or more genes have a significantly higher ncER score compared to DAEs that interact with only 1 gene (**Figure 2C**). This might reflect their biological function regulating multiple genes, resulting in a higher tendency to be constraint. A similar trend was observed for GC content, where DAEs interacting with more than one gene had a significantly higher GC content, whereas for the phastcons score, an indicator of multi-species conservation, differences were not significant (**Figure 2C**). Similar observations were made for nDAEs (**Figure 2D**). Higher GC content has also been observed in more broadly active enhancers in the immune system [45] and might be explained by binding of broadly active transcription factors (TFs) to GC-rich motifs [46].

### Sequence characteristics distinguish DAEs from nDAEs

Given the differences in gene ontology between DAE and nDAE linked genes (**Figure 1B**) and the differences in ncER score and CG content between enhancers that regulate single versus multiple genes (**Figure 2C, D**), we next asked whether there are differences between these scores in DAEs and nDAEs, and whether any potential difference would be influenced by gene interactions that these regulatory elements have. We observed a significantly higher ncER score, CG content and phastcons score when comparing all DAEs to nDAEs (**Figure 2E**). Interestingly, these scores further increased, when only considering those DAEs and nDAEs that interact with target genes (as determined by HiC). This increased even further when only considering interacting target genes that are associated with known *Online Mendelian Inheritance in Man* (OMIM) phenotypes. Similar observations were made when using the Orion [47] and CADD scores [48] (**Figure 2E**) that similarly take depletion of variation in the human population and likelihood of deleteriousness of a given nucleotide based on integration of various annotations into account, respectively. Again DAEs scored significantly higher for Orion and CADD scores than nDAEs, emphasizing the potentially biological important role of DAEs during brain development. Genes that are essential in humans are generally depleted of loss-of-function alleles, and this is reflected by a higher probability of loss-of-function intolerance (pLI) score [49]. When we plotted the median pLI of genes linked to DAEs, or to nDAEs, genes linked to DAEs scored significantly higher (**Figure 2F**). Finally, a recent study determined loss-of-function tolerance scores for non-coding sequences, by using machine learning and structural variants from whole genome sequencing, including homozygous enhancer deletions [50]. Using this analysis, we observed that DAEs were more likely to be intolerant to loss-of-function, whereas nDAEs were more often tolerant to loss-of-function (**Figure 2G**). Again, when only considering those interactions linked to known target genes, scores further improved, in favor of DAEs.

We and others previously showed that functional enhancers can be enriched for transposable elements (TEs), some of which can be human specific [51–54]. We thus asked whether DAEs and nDAEs showed a similar TE enrichment, and whether any TEs could distinguish both groups (**Figure 2H**). nDAEs showed a small enrichment for various LTR-containing TEs (e.g., LTR75B, LTR60, LTR36). Compared to nDAEs, DAEs were mainly enriched for CG rich repeat sequences, and a number of LTR repeats, such as Harlequin-int, HERVS71-int and HERVK3-int. Enrichment of the latter LTR repeats was not seen when only considering gene-interacting DAEs. The MER130 repeat family was previously shown to be enriched near critical genes for the development of the mouse neocortex and suggested to be co-opted for developmental enhancers of these genes [55]. Interestingly, MER130 repeats were enriched in all DAEs, but this enrichment was lost when only assessing DAEs that interact with genes, which made it difficult to further investigate the role of MER130 in human brain regulation. Compared to our previous findings in human embryonic stem cells (ESCs) [51], the overal TE enrichment in enhancers in brain was markedly different, with none of the TEs enriched in active enhancers in ESC showing enrichment at brain enhancers. This could indicate that different TEs co-opted into the regulatory landscape acquired different tissue specific roles during evolution.

Together this indicates that by investigating unbiased variability in epigenome marks over putative brain enhancers across developmental time points, DAEs and nDAEs can be identified which are associated with different gene ontologies, show different enrichments, have different sequence characteristics and are distinctively linked to disease relevant genes.

### DAEs and nDAEs are enriched for distinct transcription factor binding sites

As the merging of pCRs and subsequent variability calling identified DAEs with distinct sequence characteristics, we next wondered whether we could further zoom in into each of the DAEs, to identify functional relevant nucleotides. To this end, we again made use of the ncER, CG and phastcons scores, assuming that the functional relevant nucleotides in each DAE might be those that have higher scores. As the identified DAEs varied in size between 50 and 1000 bps, we first split up each DAE into 10 bp bins, and assigned the median ncER, CG and phastcons scores to each bin. To be able to compare the score distribution within each bin between all DAEs, we re-scaled each DAE to a relative bin position from 1-100 (see methods for details). Strikingly, the medians of ncER, CG and phastcons scores were highest between bins 40-60 (**Figure 3A**). To exclude that this was an artefact from the bin-rescaling, we plotted the median distribution for the same scores also for DAEs that had an identical length, and found similar results (**Supplementary Figure 3E**). We next calculated the number of reads from all epigenome data sets, and plotted the log2 enrichment over the same relative DAE bin positions. We found that ATAC-seq, DNase-seq, H3K27ac, H3K4me1 and H3K4me2 signals (all associated with enhancers) again were most enriched between bins 40-60, whereas signal for H3K4me3 (of which high levels are associated with promoters and lower levels are found at enhancers) and H3K27me3 (associated with repressed chromatin) showed a more broad distribution (**Figure 3B**), and this holds true for all developmental time points assessed. This suggests that on average the center of the DAEs most likely contains the functional relevant sequences, and given the increased chromatin accessibility at those locations, this could indicate binding of functionally relevant TFs in these central regions.

**Figure 3:**
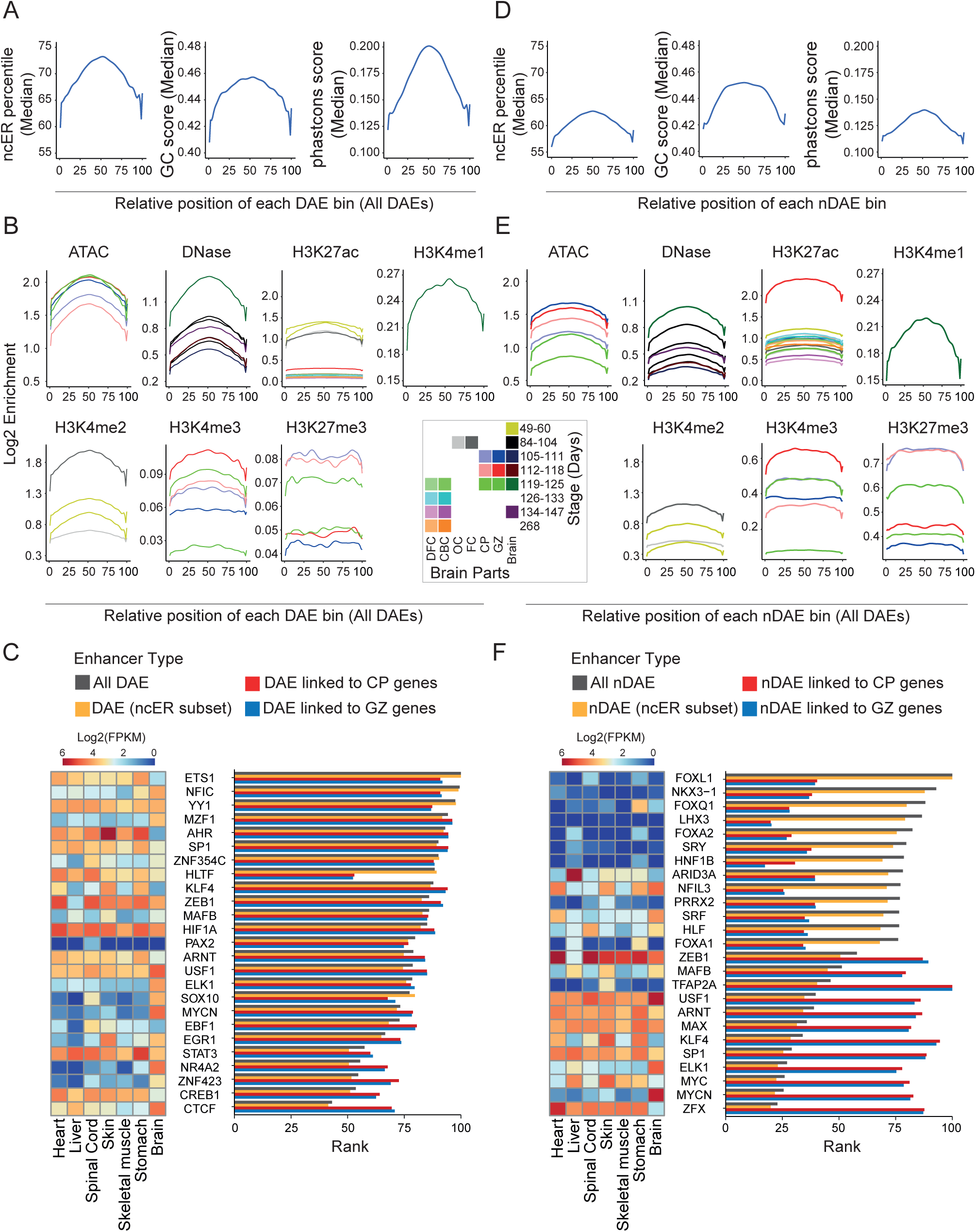
DAEs and nDAEs are enriched for distinct transcription factor binding sites. A) Line plot showing the distribution of the median ncER percentile (left) [43], GC content score (middle) [44] and phastcons score (right) [44] over the relative bin position for all DAEs. B) Line plot showing the log2 enrichment for various epigenome features as indicated, over the relative bin positions for all DAEs. Different colors indicate different time points of human brain development and different brain regions from which the data were obtained. DFC, dorsal frontal cortex; CBC, cerebellar cortex; OC, occipital cortex; FC, frontal cortex; CP, cortical plate; GZ, germinal zone; Brain, whole brain. C) Bar chart showing the relative LOLA enrichment of TFs from JASPAR in all DAEs (light gray), in the central part of all DAEs (ncER subset, orange), in DAEs linked to genes in CP (red) and in DAEs linked to genes in GZ (blue). X-axis displays the rank score (a combination of *p-*value, odds ratio from Fisher’s exact test and the raw number of overlapping regions) from LOLA. Also shown is a heatmap showing the RNA-seq expression levels (Log2 FPKM) of the same TFs across various human fetal tissues. RNA-seq data obtained from ENCODE project [7]. D) As in A), but here for nDAEs. E) As in B), but here for nDAEs. Note the difference in y-axis scale for H3K4me3 and H3K27me3 compared to panel B) given the higher enrichment in nDAEs. F) As in C), but now for nDAEs.

To investigate this further, we performed motif enrichment analysis using LOLA [56], on both full length DAEs, as well as on only the central DAE parts between bin 40-60 (ncER subset). We found a similar enrichment of TF binding sites between full length and central parts of DAEs (**Figure 3C**), and between all DAEs and those interacting with target genes in CP and GZ. This indicates that TF binding likely occurs in the central part of DAEs, and is likely implicated in the regulation of the DAE interacting genes. The most enriched TFs, included well-known TFs with essential roles for brain development. This includes amongst others ETS1, a widely studied TF with functions in different biological systems which was previously shown to be necessary for radial glia formation in vertebrates [57] and FGF-dependent patterning of anterior-posterior compartments in the *Ciona* central nervous system [58]; YY1, a crucial TF which is involved in both gene activation and repression [59], mediating enhancer-promoter interactions [60] and of which mutations cause a neurodevelopmental disorder [61]; and CTCF, a master regulator of chromatin structure, of which *de novo* mutations cause intellectual disability [62]. We next repeated the same analysis for nDAEs (**Figure 3D-F**). Similar to our observations for DAEs, nDAEs had higher ncERs, CG content and conservation at the central part, with those regions being enriched for enhancer associated histone marks, but showing less variability over time. When performing TF enrichment analysis, we observed a different and less specific set of TFs enriched at nDAEs compared to DAEs. Also, enrichment was lower at those nDAEs that were interacting with target genes. Again similar enrichment was found in the central part compared to the whole nDAEs, indicating that TF binding sites tend to be located in the middle of these enhancers. Enriched TFs for nDAEs included amongst others FOXL1, a transcriptional repressor that regulates central nervous system development [63]; the LIM homeodomain TF LHX3, that is essential for pituitary and nervous system development [64, 65]; and FOXA2, which plays a role in midbrain dopaminergic neurons [66, 67] (**Figure 3F**). Shared TFs enriched both at DAEs and nDAEs included SP1, which loss in astrocytes impacts on neurons in the cortex and hippocampus of mice [68]; MAFB, a basic leucine zipper TF that plays a role in hindbrain development [69–71] and postnatal brain development [72, 73]; and ZEB1 which is required for neuronal differentiation [74, 75].

Together this indicates that on average the central part of brain enhancers (both DAE and nDAEs) contains relevant but partially distinct TF binding sites and might be enriched for functional relevant sequences, which can be further fine-mapped using ncER scores and other sequence characteristics. To test this directly, we selected three DAEs, linked to *IRF2BPL*, *CHD2* and *MACF1*, that showed activity in reporter assays in NSCs (**Figure 1G**), and deleted 10-30 bp of those regions that had the highest ncER scores in those enhancers. Upon transfection of these mutant DAEs, we observed a significantly reduced enhancer activity for *IRF2BPL* and *CHD2*, but not for *MACF1* (**Supplemental Figure 4**). Deleting regions with a lower ncER score did not affect enhancer activity. Together this indicates that integrative analysis, variability analysis during development and sequence characteristics can identify functional relevant nucleotides in brain enhancers.

### Epigenome at DAEs show temporal dynamics during human brain development

To further understand the dynamics of enhancer regulation, we subdivided DAEs interacting with genes in GZ and CP by performing clustering analysis on all available epigenome data sets, at different developmental stages (between 8-12 PCW, 13-18 PCW and >18 PCW) (**Figure 4A**). At 8-12 PCW, we found two clusters for both GZ and CP that showed relatively constant enrichments over time, with the first cluster (red) showing a higher enrichment for all epigenome features available for that developmental stage, compared to the second cluster (green). No statistical significant differences in gene expression levels between genes linked to both clusters were found. Genes associated with cluster 1 DAEs in CP were enriched for gene ontology terms related to neuronal differentiation, whereas cluster 2 was dominated by processes in the Golgi. Likewise, for GZ, genes associated with cluster 1 seemed to be associated with more specific biological functions, whereas processes associated with cluster 2 showed more broad involvements (**Figure 4A**).

**Figure 4:**
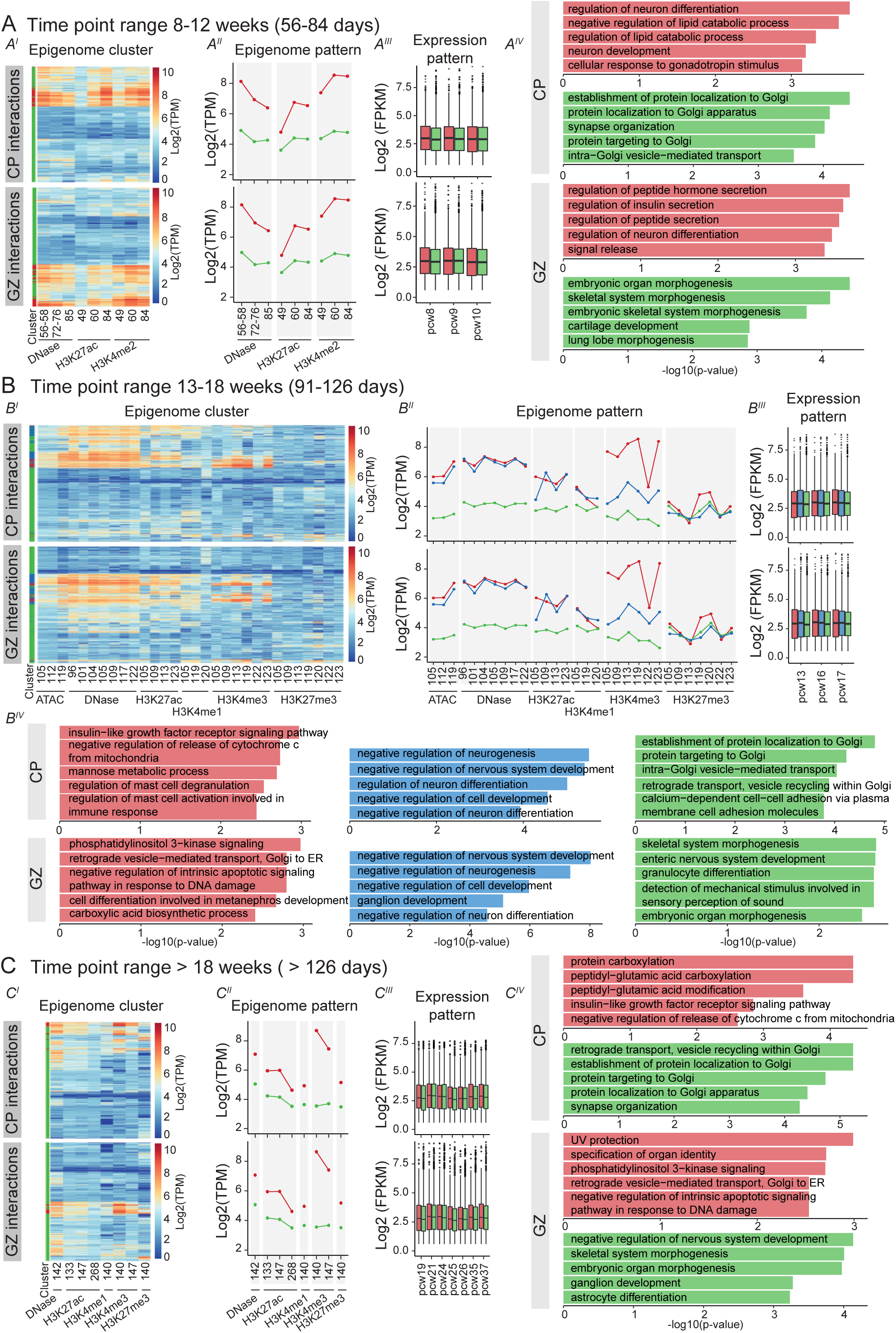
Clustering of DAEs unravels temporal dynamics of brain gene regulation. A) Heatmap displaying all available epigenome features for PCW 8-12, across all DAEs interacting with protein coding genes in CP (upper heatmap) and GZ (lower heatmap) (A^I^). K means clustering analysis of epigenome features (A^II^) identifies two clusters, cluster 1 (red) and cluster 2 (green). Level of enrichment is indicated on the y-axis in Log2 TPM. Box plots (A^III^) shows RNA-seq gene expression of protein coding genes regulated by the DAEs from each cluster (Expression pattern), for available data from PCW 8, 9 and 10 [106]. Boxes are IQR; line is median; and whiskers extend to 1.5 the IQR. Gene enrichment analysis for the corresponding genes in each cluster (A^IV^). X-axis shows the −log 10 (p-value) from Enrichr. B) As for A), but now for PCW 13-18. C) As for A), but now for PCW >18.

At 13-18 PCW, three clusters emerged in both GZ and CP (**Figure 4B**). Whereas cluster 3 (green) showed relatively low levels of epigenome marks similar to cluster 2 at 8-12 PCW, cluster 1 (red) and cluster 2 (blue) showed higher epigenome enrichments. Both cluster 1 and 2 had similar levels of H3K27ac, but mainly diverged from each other on the levels of H3K4me3. Cluster 2 was strongly enriched for processes involved in neural system development both in CP and GZ. Gene ontology of genes associated with cluster 1 (red) which showed higher H3K4me3 levels, showed enrichment for insulin-like growth factor receptor signalling and immune cell related processes in CP. Insulin-like growth factors are important for neuronal survival and neurogenesis [76]. As high levels of H3K4me3 have also been found at enhancers in blood cells [77], possibly stabilizing their transcription, it is tempting to speculate that part of this cluster reflects enhancers active in hematopoietic cells from the developing vasculature [78] and microglia (brain tissue macrophages) that are invading the brain at these developmental time points [79]. In GZ, cluster 1 was associated with phosphatidylinositol 3−kinase signaling, which is important for commitment of neural progenitor cells [80, 81].

Finally, at >18 PCWs, we found two clusters of DAEs, of which cluster 1 (red) was marked by higher levels of epigenome marks. (**Figure 4C**). In CP, genes associated with this cluster were enriched for carboxylation processes and insulin-like growth factor receptor signalling. Genes associated with the second cluster (green) were again more enriched for broad developmental processes, including the Golgi system. In GZ, genes associated with cluster 1 (red) were amongst others involved in DNA damage repair. Indeed, alterations in this pathway can lead to reduced proliferation of neural progenitor cells leading to microcephaly [82, 83]. Cluster 2 (green) in GZ was associated with terms related to neurodevelopment and organ morphogenesis.

Together, this shows that temporal epigenomic rearrangement in DAEs is reflected in regulating the expression level of genes that are important in developmental and cell type specific processes.

### DAEs regulate disease relevant genes and are enriched for disease implicated variants

Given our findings that DAEs are associated with genes relevant for brain development, we further investigated which disease relevant genes are regulated by DAEs. We first focused on known disease causing genes retrieved from OMIM. We found that 1,556 OMIM genes are regulated by DAEs (of which 1,165 and 1,166 from the interactions found in GZ and CP, respectively). Most DAEs are linked to genes involved in mental retardation, developmental and epileptic encephalopathy, and neurodevelopmental disorders (**Figure 5A**). This included genes like *KMT2C*, involved in Kleefstra syndrome (OMIM #617768), and *GRIN2A* of which heterozygous mutations cause epilepsy and speech delay (OMIM #245570). Next to genes, enhancers can also interact with other additional enhancers. Interestingly, the more additional enhancers (DAE and/or nDAE) a DAE was interacting with, the more likely the target gene of this DAE was an OMIM gene (**Supplementary Figure 3F**). This supports recent findings that the number of enhancers linked to a gene reflect its disease pathogenicity [84], and confirms enhancer redundancy for disease relevant genes [85].

**Figure 5:**
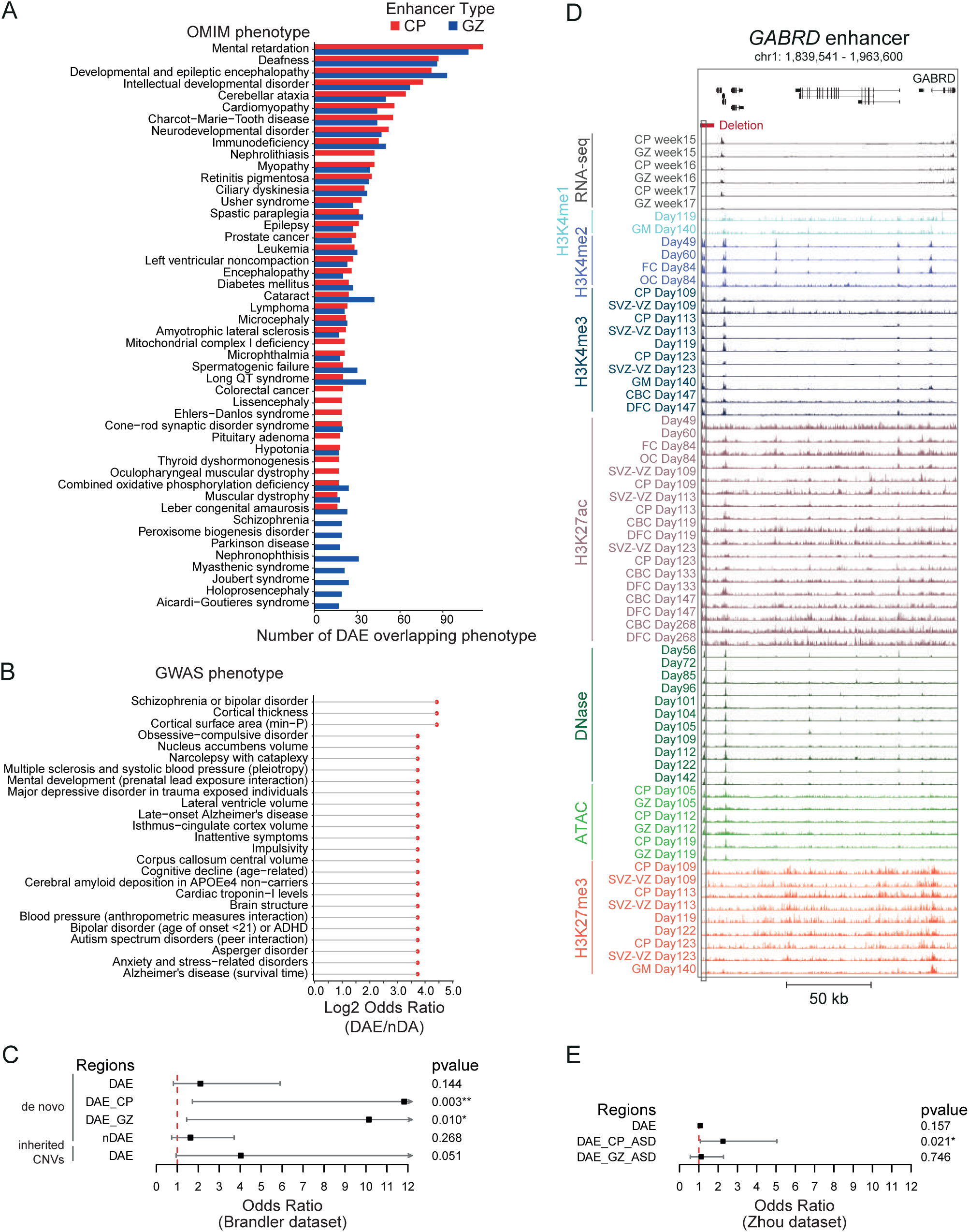
Variants in DAEs and nDAEs are associated with human disease. A) Bar graph showing the number of DAEs linked to their target genes in CP and GZ and their most enriched OMIM phenotypes. B) Plot showing the top-25 GWAS phenotypes that are enriched in DAEs compared to nDAEs (log2 odds ratio DAE/nDAE). C) Line graph showing the odds ratio, confidence interval and *p*-value for enrichment of CNVs from an ASD cohort at DAEs and nDAEs. CNVs data obtained from Brandler et al [86]. D) Line graph showing the odds ratio, confidence interval and *p*-value for enrichment of SNV from an ASD cohort at DAEs and nDAEs. SNV data obtained from Zhou et al [88]. E) Genome browser track showing the regulatory landscape of the *GABRD* gene. Indicated are a DAE (chr1: 1,840,449-1,840,835) that is interacting with the *GABRD* promoter, and a deletion (chr1: 1,840,001-1,845,000) that is found in an epilepsy patient (CNET0068) from Monlong et al [87]

We next leveraged published GWAS loci for brain-related traits and disorders. When comparing the odds ratio between DAEs and nDAEs, we found that DAEs were more often enriched for various significant GWAS loci, reflecting a broad variety of both brain developmental processes (e.g., volumes of different anatomical brain regions) and neurodevelopmental disorders (e.g., mental development, autism) (**Figure 5B**).

Encouraged by these findings, we next asked whether copy number variants (CNVs) or single nucleotide variants (SNVs) at DAEs could be involved in causing genetic disease. We first leveraged previously published disease implicated CNVs. Brandler et al performed WGS in their discovery cohort of individuals affected by an autism spectrum disorder (ASD) and unaffected individuals, and reported on 135 *de novo* CNVs (104 deletions, 29 duplications and 2 inversions) [86]. Of these, 25 overlapped a DAE in cases, and 8 in controls (odds ratio=2.10, *p*-value=0.144101). When only considering those CNVs overlapping DAEs linked to target genes, this became 17 in cases and 1 in control for DAEs linked to CP genes (odds ratio=11.83, p=0.003003) and 15 in cases and 1 in control for DAEs linked to GZ genes (odds ratio=10.14, *p*-value=0.010423). For nDAEs, 36 CNVs were found in cases and 15 in controls (odds ratio=1.63, *p*-value=0.267964). However, as not all these CNVs exclusively covered non-coding regions, it cannot be excluded that the observed association is due to disrupted coding genes, rather than involvement of DAEs. We therefore also assessed rare inherited deletions from the same study that did not overlap with coding exons (n=213 in total, 175 in cases and 38 in controls). From these, 32 cases had a deletion covering a DAE, compared to two controls (odds ratio=4.027972, *p*-value=0.05119). Although not significant, this might point to more deletions covering DAEs in ASD individuals, but would require a larger sample size to be confirmed (**Figure 5C**).

In another study, Monlong et al [87] reported on CNVs in 198 epilepsy patients detected by WGS. They found an enrichment of rare non-coding CNVs near known epilepsy genes, with the *GABRD* gene showing the strongest and only nominally significant association with 4 non-coding deletions amongst the epilepsy patients. Interestingly, a 4999 bp deletion reported in that study, overlapped with a 386 bp DAE which is located ∼110 kb upstream of *GABRD* and which interacts with its promoter (**Figure 5D)**. Hence it is possible that deletion of this DAE affects *GABRD* expression, which might be implicated in the phenotype of that individual.

Third, we made use of *de novo* SNVs found in WGS from 1,790 ASD simplex families [88]. We found 932 *de novo* variants that overlapped all DAEs in ASD individuals compared to 829 variants overlapping all DAEs in unaffected individuals (odds ratio=1.07, *p*-value=0.157). We next repeated the analysis with only those DAEs that are interacting with known autism genes from the SFARI Gene database (n=1,003 genes) [89]. We found 26 cases and 11 controls with *de novo* variants in DAEs that interact with autism genes in CP (odds ratio=2.249703, *p*-value=0.021455), whereas for DAEs interacting with autism genes in GZ this was 20 cases and 17 controls (odds ratio=1.11955, *p*-value=0.745628) (**Figure 5E**). Interestingly, for the genes *CIB2*, *FBRSL1*, *PACS2*, *KDM4B* and *MYT1L* we each found 2 individuals with autism with *de novo* variants in DAEs interacting with these genes. These variants are either absent or extremely rare in a large control cohort of gnomAD [90], possibly pointing to a role in causing the phenotype, although this will require further validation.

Together this indicates that DAEs are linked to disease relevant genes, and are enriched for GWAS loci relevant for brain related traits and for variants linked to genetic disorders.

### CRISPRi and zebrafish experiments confirm enhancer activity of DAEs regulating genes involved in epileptic encephalopathy

To further substantiate our findings, we validated the biological role of selected enhancers, using *in vivo* zebrafish transgenic reporter assays and CRISPR inhibition in human NSCs by focussing on enhancers linked to disease relevant genes.

*CHD2* belongs to the chromodomain helicase DNA binding families of chromatin remodeling proteins, and haplo-insufficiency of this gene has been associated with a developmental and epileptic encephalopathy, presenting with early onset intractable seizures, cognitive regression, intellectual disability and ASD behaviors (OMIM #615369) [91]. Around 80 kb upstream of *CHD2*, we found a DAE that interacts with the *CHD2* promoter (**Figure 6A**). In NSC reporter assays, this region showed strong enhancer activity and this was less pronounced in non-neural HEK cells (**Figure 1G**). To further study the biological relevance of this region, we first tested enhancer activity *in vivo* using zebrafish transgenesis. Out of the 36 analyzed zebrafish larvae, 61.1% showed GFP-expression in the forebrain at 1 day post fertilization (dpf), and this increased to 81.8% at 2dpf and 87.9% at 3dpf, indicating enhancer activity (**Figure 6B,C**). Expression was also found in midbrain and hindbrain, at a slightly lower extent, in the eyes, in peripheral neurons and in the spinal cord. GFP expression in the developing zebrafish brain correlated with *in situ* hybridisations of endogenous chd2 (https://zfin.org/ZDB-FIG-060810-2123) [92]. To test whether epigenome silencing of this enhancer would affect *CHD2* expression, we performed CRISPR interference (CRISPRi) by targeting dCas9-KRAB-MeCP2 to the enhancer region by co-expression of gRNAs with a GFP fluorescent reporter. Transfection efficiency in these experiments, based on FACS for GFP, was 78-92%, and this resulted in around 50% reduction of *CHD2* expression compared to mock cells transfected solely with dCas9-KRAB-MeCP2 (**Figure 6D**). Interestingly, it was previously shown that silencing of *CHD2* leads to reduced expression of *REST* [93]. In agreement with this, cells with reduced *CHD2* expression upon *CHD2* enhancer silencing showed reduced *REST* expression (**Figure 6E**). This confirms that *CHD2* is under control of the investigated DAE.

**Figure 6:**
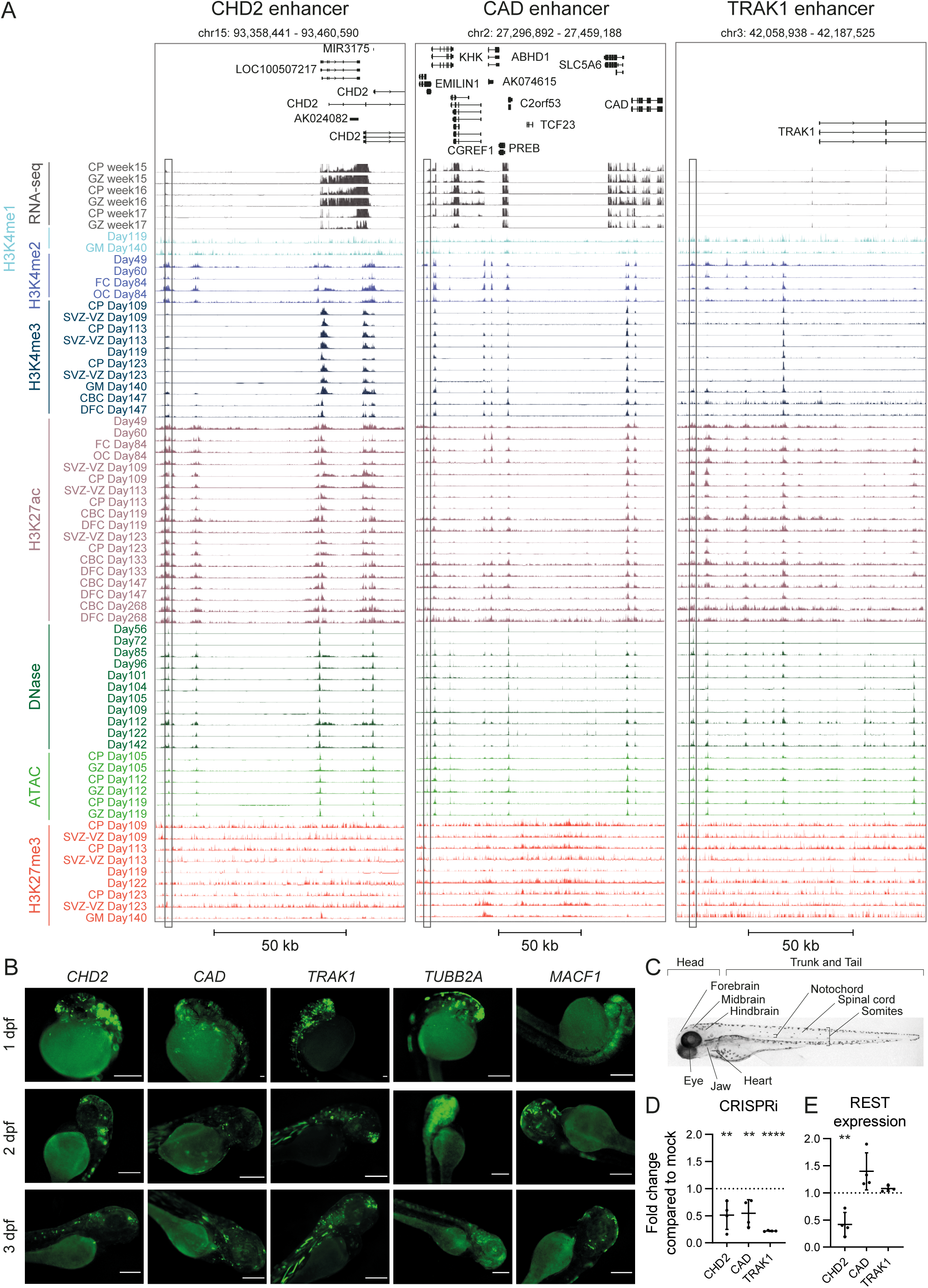
CRISPRi and zebrafish experiments validate activity of DAEs regulating genes involved in neurogenetic disorders. A) Genome browser tracks showing enhancers interacting with *CHD2* (left), *CAD* (middle) and *TRAK1* (right). Shown are RNA-seq expression profiles, various histone modifications, and ATAC-seq and DNase profiles for various time points during human fetal brain development, as indicated. The tested DAEs are indicated by the box. B) Representative fluorescent images showing GFP expression of transgenic enhancer reporter assays in zebrafish larvae at 1, 2 and 3 dpf. Tested are the enhancers for *CHD2*, *CAD* and *TRAK1* (shown in A), and two additional enhancers for *MACF1* and *TUBB2A*. The five tested enhancers induced GFP-expression in the head of the larvae, amongst others in the forebrain in 61.1%, 81.8% and 87.9% larvae for *CHD2*; 88.9%, 85.4% and 85.7% for *CAD*; 87.1%, 70% and 88.5% for *TRAK1*; 81.5%, 85.7% and 76.2% for *MACF1*; and 87.5%, 100% and 100% for *TUBB2A*, respectively at 1, 2 and 3 dpf. Also peripheral neuron-specific GFP expression was found, with 0%, 60.6% and 21.2% for *CHD2*; 68.9%, 24.4%, and 51.4% for *CAD;* 83.6%, 65.5% and 67.3% for *TRAK1*; 37%, 50% and 33.3% for *MACF1*; and 50%, 83.3% and 63.3% for *TUBB2A*, respectively at 1, 2 and 3 dpf. Scale bars represent 500 µm. C) Bright field image of a wild type zebrafish larvae at 3dpf (lateral view), with the anatomical sites that were scored for GFP expression indicated. D) qRT-PCR showing reduction of *CHD2*, *CAD* and *TRAK1* expression in NSCs upon silencing of respective enhancer by dCas9-KRAB-MECP2. Data represent fold change of expression of respective genes compared to mock transfected cells (KRAB-MECP2 plasmid only, no gRNA plasmid). Two independent transfection experiments were performed, each in duplicate. All data points and standard deviation are shown. ** p<0.01; *** p<0.001 (one-way ANOVA test followed by multiple comparison test (Fisher’s LSD test) E) qRT-PCR showing reduction of *REST* expression in NSCs upon silencing of *CHD2*, *CAD* or *TRAK1* enhancers by dCas9-KRAB-MECP2. Data represent fold change of REST expression compared to mock transfected cells (KRAB-MECP2 plasmid only, no gRNA plasmid). Two independent transfection experiments were performed, each in duplicate. All data points and standard deviation are shown. ** p<0.01; *** p<0.001 (one-way ANOVA test followed by multiple comparison test (Fisher’s LSD test).

Bi-allelic variants in *CAD* cause an early infantile epileptic encephalopathy (OMIM #616457) [94], that is characterized by global developmental delay, loss of skills, therapy refractory epilepsy, brain atrophy, and dyserythropoietic anemia. We found an enhancer located in the third intron of *EMILIN1*, around 135 kb upstream of *CAD*, that interacts with the *CAD* promoter (**Figure 6A**) and which showed strong enhancer reporter activity in NSCs and only limited activity in HEK cells (**Figure 1G**). Targeting this region in NSCs by CRISPRi significantly diminished gene expression of *CAD* to around 50% compared to mock (**Figure 6D**). Similar to *CHD2*, *in vivo* reporter assays in zebrafish recapitulated *in situ* hybridisation results for cad (https://zfin.org/action/figure/all-figure-view/ZDB-PUB-010810-1?probeZdbID=ZDB-EST-021105-21) [95]. From the 45 analyzed larvae, GFP expression was found in the forebrain of 88.9% larvae at 1dpf, which remained ∼85% at 2 and 3dpf. Again, GFP expression was observed also in midbrain, hindbrain, eyes, in peripheral neurons, notochord and spinal cord (**Figure 6B**).

We next focused on an enhancer interacting with *TRAK1*, located ∼65 kb upstream of the TSS (**Figure 6A**). *TRAK1* is involved in mitochondrial trafficking, and bi-allelic loss-of-function variants in *TRAK1* are associated with developmental and epileptic encephalopathy (OMIM #618201) [96, 97]. Similar to the *CHD2* enhancer results, the *TRAK1* enhancer showed higher reporter assay activity in NSCs than in HEK cells (**Figure 1G**). Targeting of dCas9-KRAB-MeCP2 to the *TRAK1* enhancer reduced *TRAK1* expression to ∼25% residual expression (**Figure 6D**). Interestingly, in the VISTA enhancer browser, another enhancer linked to *TRAK1* (hs2359), ∼18 kb upstream of the TSS, has been reported which did not show enhancer reporter activity in E11.5 mouse embryos. When testing the *TRAK1* enhancer identified here in zebrafish (**Figure 6B**), we found that from 55 larvae, 89.1% showed GFP-expression in the forebrain, as well as in the midbrain (74.5%) and hindbrain (85.5%). The larvae showed decreasing GFP expression in neurons outside of the brain over the different time-point (83.6% at 1dpf, 65.5% at 2dpf and 67.3% at 3dpf) and increasing expression in both the somites (89.1%) and the heart (58.2%) at 3dpf, compared to 32.7% and 1.8% at 1dpf larvae, respectively. Moreover, this enhancer was active also in the eye, trunk and tail, notochord and, at 1 dpf, in the spinal cord.

Finally, next to these three enhancers, we validated 7 additional enhancers linked to the genes *LRP1*, *LRP5*, *TUBB2A*, *ELOVL6*, *MACF1*, *C12orf4*, and *EBP41L1* using zebrafish reporter assays, and could confirm enhancer activity for all of them with >60% larvae expressing GFP (**Figure 6C**, **Supplementary Figure 5**). These included enhancers linked to the disease genes *MACF1* (OMIM #618325) and *TUBB2A* (OMIM #615763), of which coding pathogenic mutations cause brain malformations [98, 99], and *C12orf4* (OMIM #618221) of which bi-allelic variants cause intellectual disability [100]. Together, this shows that DAEs identified in this integrative analysis show enhancer activity *in vitro* and *in vivo* and regulate amongst others genes linked to Mendelian disorders.

## Discussion

Here we perform an integrative analysis of virtually all previously described putative enhancers and epigenome data sets of relevance for human brain development. We identify almost 40 thousand enhancers that display epigenomic rearrangement during brain development. These DAEs have different sequence characteristics compared to non-variable enhancers, are bound by distinct sets of TFs, regulate disease relevant genes and can harbor non-coding variants that are associated with human disease.

Our integrative analysis identified a large number of enhancers linked to known disease genes, and expands on the knowledge of regulation of these genes. For example, *CHD2* expression regulation has so far only been known to be influenced by a highly conserved long non-coding RNA (lncRNA) referred to as CHD2 Adjacent Suppressive Regulatory RNA, CHASERR), which is located in proximity to the *CHD2* TSS, and which represses *Chd2* gene expression *in cis* [101]. It has been hypothesized that targeting CHASERR could be used to increase expression of *CHD2* in haploinsufficient individuals [101], and it will be interesting to explore whether targeting the enhancer region of *CHD2* that we find and validate here could be exploited as an alternative target of such a strategy. Similarly, the regulation by enhancers of the other disease implicated genes that we validate here adds to the list of potential targets to find disease causing non-coding variants that disturb this regulation.

An interesting finding of our study is that by starting with putative enhancers and variability of epigenome features over time during development, we recover DAEs and nDAEs that can be distinguished based on sequence characteristics, such as differences in GC content, the level of sequence constraint, tolerance to loss-of-function and differential profiles of TF binding. Also, these DAEs and nDAEs seem to be associated with distinct developmental processes, and result in differences in gene expression levels. It is tempting to speculate that the distinctive features between these two types of enhancers can be used to uncover key nucleotides responsible for those biological regulatory differences. It seems plausible that disturbing of these functionally causative sequences could lead to altered physiology resulting in disease. Our analysis revealing GWAS loci enrichment and the link of DAEs supports this statement. We suggest that our results might help interpreting the effects of SNVs in non-coding sequences, which is at this stage not a trivial task. Our annotated database of DAE and nDAE will be instrumental to prioritize SNVs based on distinct sequence characteristics identified for these elements as well as to provide cues on potentially disturbed developmental processes based on differential temporal activity and regulatory targets of the enhancer in question. This in turn can instruct functional validation and help deciphering pathogenicity of variants. With an increasing number of whole genome sequencing data available, it is expected that more, possibly disease implicated, non-coding variants will be identified, and the need to classify those sequences in benign or pathogenic will only further increase. With more computational pathogenicity prediction tools available, such as the ncER score and outcomes of integrative analyses such as performed here that pinpoint likely functional sequences, it might become possible to further decipher the impact of these SNVs.

## Conclusions

We foresee that this comprehensive compendium of (likely) functional enhancer sequences during brain development will be instrumental to identify disease causing variants which might explain parts of the missing heritability in the field of clinical genetics.

## Methods

### Data collection and processing

#### Collection of putative brain enhancers

To generate a comprehensive set of putative brain enhancers active during fetal brain development, we scrutinized Pubmed and various enhancer databases (last assessed: April 2019), including amongst others EnhancerAtlas, Fantom project, Vista Enhancer. This resulted in 1,595,292 putative enhancers. Enhancers with identical coordinates were deduplicated and the unique regions were used to determine putative critical regions (pCRs), reasoning that overlapping parts of a putative enhancer obtained from different sources might point to functional relevant regions of that putative enhancer. If there is any overlap between coordinates of putative enhancers derived from two or more databases, the pCRs were defined as the overlapping regions present in those databases using BEDtools (mergeBed, intersectBed, genomeCoverageBed and groupBy commands) (version 2.30.0) [102]. Putative enhancers that were only present in one of input sources were also included in the pCRs (**Figure 1, step 1**), as we cannot exclude that these putative enhancers are biologically relevant. pCRs with length less than 50bp and more than 1000bp were excluded. To avoid any overlap with promoters, enhancers located within 2kb upstream or 1kb downstream of a transcriptional start site (TSS) (Ensembl GRCh37.p13 Release 102) were excluded using intersectBed(version). Following this procedure we identified a total of 202,462 pCRs which were used for downstream analyses. Next, we excluded 299 pCRs that were not covered by sufficient amounts of epigenome data (less than 10 reads in (see section on defining DAEs), resulting in a final number of 202,163 pCRs. GREAT web interface was used (version 4.0.4) (http://great.stanford.edu/public/html/)[36] to visualize enhancer-TSS distance (with *basal plus extension, proximal 5kb upstream and 1kb downstream, plus distal up to 100kb, including curated regulatory domains, and whole genome (GRCh37/hg19) as background* parameters) (**Supplementary Figure 2B**).

#### Epigenome data

Epigenome data was collected from Roadmap Epigenomics Consortium, ENCODE, PsychENCODE and other studies. Epigenome data sets used for integration included histone modifications (H3K27ac, H3K27me3, H3K4me1, H3K4me2, H3K4me3) and chromatin accessibility (ATAC-seq and DNase-seq) from different brain regions and different human developmental stages (**Figure 1,step 1**). To avoid any possible confounding biases because of the various pipelines used in different studies, we reanalyzed the raw FASTQ files using our analysis pipeline (**Supplementary Figure 1**). First, adaptor contamination was removed using Trim Galore (version 0.6.5 (https://www.bioinformatics.babraham.ac.uk/projects/trim_galore/) and trimmed data were aligned to the GRCh37/hg19 human genome using Bowtie2 aligner (version 2.4.2)(with *--very-sensitive* parameter) [103]. Only properly paired and uniquely mapped reads, with mapping quality more than 30 (MAPQ >=30), were kept followed by removing any possible duplicated reads using Picard’s MarkDuplicates (version 4.0.1.1)(http://broadinstitute.github.io/picard/). These reads were used to define differentially active enhancers (DAEs).

### Defining differentially active enhancers (DAEs)

We assumed that pCRs with high variability in different epigenome data (dynamic epigenomic rearrangement) across different developmental stages are more likely to be functional than other pCRs. To determine this variability, the number of overlapping reads (for each epigenome mark) with pCRs were counted using the multiBamCov command of BEDtools and a matrix was generated that included enhancers as rows and epigenome features as columns. Epigenome features were from different brain regions and developmental stages. 299 pCRs with less than 10 reads were excluded, leaving 202,163 pCRs for this analysis. Subsequently, the raw read count matrix was normalised using TMM-normalization [104] and the normalized count matrix was used to define DAEs across different developmental stages and brain regions of a given epigenome data using edgeR (version 3.32.1)[35]. Since there were different developmental stages (time-point factor) and brain regions (brain part factor) in each epigenome data, a design matrix was generated for each factor separately. A limited number of samples without biological replicates were grouped together with other samples based on high correlation (Pearson correlation; *r* > 0.89). The DAEs were defined based on each design matrix using a generalized linear model and quasi-likelihood F-tests. In order to define the final DAE list, DAEs identified from at least two epigenome data specific matrices were pooled. In total, this resulted in 39,709 DAEs (FDR adjusted *p-value* < 0.05). The remaining 162,454 pCRs that did not show variability were considered as nDAEs.

### Identifying chromatin interactions

#### Enhancer-Gene interactions

In order to define Enhancer-Gene interaction, published HiC data from 3 human fetal brains, for cortical plate (CP) and germinal zone (GZ) at gestation week 17–18 were used [26]. This data provides 10kb resolution bins for gene loop interactions and 40kb resolution for topologically associating domain (TAD). Pre-calculated significant interactions were intersected with pCRs (DAEs and nDAEs) using intersectBed to define gene-enhancer interaction for both CP and GZ separately. Out of the 202,163 pCRs, 41,041 pCRs engaged in 101,366 interactions in CP, and 41,085 pCRs had 100,521 interactions in GZ. Enhancer-gene interactions locating within the same TAD were considered for downstream analyses (almost 80% of all interactions were intra-TAD). We only included protein coding and lincRNA genes in our analysis. To determine enhancer-enhancer interactions in **Supplementary Figure 3F** we also intersected HiC data with pCRs, focussing on interactions between DAEs and both DAEs and nDAEs.

In addition to HiC, we employed other enhancer-gene interaction predictions including JEME (http://yiplab.cse.cuhk.edu.hk/jeme/) [37], ENCODE (https://ernstlab.biolchem.ucla.edu/roadmaplinking/) [38], FOCS (http://acgt.cs.tau.ac.il/focs/download.html) [39] and GeneHancer (downloaded from UCSC table browser; hg19; updated 2019) [40]. These databases apply statistical models on different types of omics data to predict enhancer-gene interactions. We collected fetal brain enhancer-gene predictions from JEME and ENCODE and all brain related enhancer-gene predictions from FOCS and GeneHancer, as the latter two resources do not specify fetal specific interactions. Intersections between the pCRs and each of these predictions were considered as enhancer-gene interaction.

### Functional enrichment analysis

#### Enhancer sequence characteristics analysis

To determine whether different DNA sequence features distinguish different enhancer groups and whether there is any association between these features and functional prediction, we considered the following features: (i) the non-coding essential regulation (ncER) score (https://github.com/TelentiLab/ncER_datasets/<u>; updated 06-03-2019)</u> [43]; (ii) GC content, as determined by the GCcontent R packages based on BSgenome.Hsapiens.UCSC.hg19 (version 1.4.3); (iii) conservation score for each enhancer, as derived from the gscores R packages based on phastCons100way.UCSC.hg19 (version 3.7.2) [44]; (iv) Orion scores [47]; (v) CADD scores [48]; (vi) Haploinsufficiency scores [50] and (vii) probability of loss-of-function intolerance (pLI) score [49]. The overlaps between DNA sequence features and enhancer coordinates were defined using intersectBed. As assessed enhancers (e.g., pCRs) varied in length between 50-1000 bp, and the above mentioned scores were given either at the nucleotide level or in certain bins (depending on the given scores from the individual resources), we calculated the median value for each enhancer and used this in group comparisons. For gene specific scores (e.g., pLI), we plotted the scores of the genes linked to the enhancers. Statistical significant differences between groups were determined using Wilcoxon signed rank test in R.

#### Gene expression correlation

To compare gene expression levels of enhancer target genes between different groups, various transcriptome data were collected. This included: transcriptome data from different brain regions and developmental stages, and also various control data from other fetal tissues from the Roadmap Epigenomics Consortium, ENCODE project, Allen human brain atlas and other studies [7, 10, 30, 105, 106]. Raw data (Fastq) was quality controlled and adaptors and other contaminants were removed using Trim Galore (version 0.6.5), reads were mapped to the GRCh37/hg19 human genome assembly using STAR aligner (version 2.7) [107], and gene counts were obtained using htseq-count (version 0.12.4) [108]. Gene expression levels were normalized based on fragments per kilobase of transcript per million mapped reads (FPKM). To correlate enhancers to gene expression, enhancer-gene interactions were derived from the HiC data as described above. Gene expression levels were plotted and statistical comparison was performed, between expression levels of subgroups, using Wilcoxon signed rank test in R.

#### Gene ontology analysis

For functional enrichment analysis, we used GREAT [36] and Enrichr [109]. GREAT was used via the web interface (version 4.0.4) (http://great.stanford.edu/public/html/) using the following settings: basal plus extension, proximal 5kb upstream and 1kb downstream, plus distal up to 100kb, including curated regulatory domains, and all pCRs as background. The −log 10 binomial *p*-value was used to rank GREAT enrichment. Enrichr was used via the web interface (https://maayanlab.cloud/Enrichr/). Enrichr was used with default parameters and the whole genome set as background.

#### Transcription factor binding enrichment

We used LOLA [56] to assess binding of known transcription factors to DAEs and nDAEs (**Figure 3**). We used motifs from the JASPAR motif database (using reference genome GRCh37/hg19 and LOLAJaspar database core), to test the motif enrichment in DAEs and nDAEs, using all pCRs as background. The combined rank index (a combination of *p-*value, odds ratio from Fisher’s exact test and the raw number of overlapping regions), was used to rank the known motifs.

#### Transposable element enrichment

The RepeatMask (GRCh37/hg19, updated 20-02-2020) was downloaded from the UCSC table browser and joined to the pCRs. To determine enrichment of transposable elements in brain enhancers, we followed a strategy previously used when investigating active enhancers in human embryonic stem cells [51]. The number of overlaps of each type of repeat (n_overlaps) with all pCRs (n) was used to calculate the relative frequency (f_all = n_overlaps/n). Multiplication of the relative frequency with the number of regions (n_test, e.g., DAE, nDAE etc.) in any tested group yields the expected frequency (E). This number was compared with the actual observed frequency in the subgroups (f_test = (n_overlap, test)/n_test = O) to calculate the observed versus expected ratio (O/E). We considered repeats with O/E < 0.5 as depleted, or O/E > 1 as enriched. For the subsequent data interpretation we only focused on transposable elements that were present multiple times (n_overlap > 15) in all pCRs.

#### Disease relevance enrichment

The *Online Mendelian inheritance in Man* (OMIM) gene list (updated 28-09-2020) was downloaded using biomaRt R package [110] from Ensembl GRCh37.p13 Release 101. The GWAS catalog (GRCh37/hg19,updated 17-03-2021) was downloaded from the UCSC table browser. The GWAS catalog was manually filtered to keep brain related studies and their variants. For CNV analysis, we retrieved pre-processed published data from Brandler et al (their supplemental table 9: de_novo_SVs sheet, and their supplemental table 7: Primary CR Trans and Replication CR Trans sheets) [86] and Monlong et al (cnvs-PopSV-Epilepsy-198affected-301controls-5kb.tsv.gz file in https://figshare.com/s/20dfdedcc4718e465185) [87]. For SNV analysis of the ASD simplex families, we collected *de novo* variants from supplemental table 1 of Zhou et al [88]. Autism genes were collected from the SFARI Gene database (http://gene.sfari.org/database/human-gene) [89]. The overlap between enhancer regions (DAE and nDAE) and each data set was determined using intersectBed. The odds ratio and *p*-value between DAE and nDAE was calculated using fisher.test () R function. The Haldane–Anscombe correction was used to adjust the odds ratio. We generated 500,000 random genomic regions with an average length of 500bp based on GRCh37/hg19 using creatRandomRegions () R function.

### Distribution of features across enhancer bins

To see the distribution of enrichment of different features (ncER score, GC content, phastCons score and epigenome data) across enhancers, we divided the enhancer regions into 10 bp bins and calculated the relative scores (the median value for ncER score, GC content, phastCons score) and the number of reads (for epigenome data) for each bin. As the enhancers under investigation differed in size between 50-1000 bp, to make enrichments between enhancers comparable, we re-scaled each enhancer bin. To this end, we calculated a relative position between 1-100 for each bin of each enhancer, where 1 is the first bin, and 100 is the last bin of each individual enhancer. We then plotted the distribution of each feature across all these re-scaled enhancer bins.

### DAE clustering analysis

The matrix of DAEs was used to determine the pattern of epigenome data through different developmental stages. To determine the optimal clustering algorithm, we used clValid R package which simultaneously compares multiple clustering algorithms (hierarchical, kmeans, model-based, pam and clara). Based on this, the pam algorithm (which is similar to k-means but more robust to noise and outliers) was selected to cluster DAEs using the spearman distance and ward.D^2^ method. To define the optimal number of clusters, we used fviz_nbclust and NbClust R packages which compute different indices by bootstrapping (n=1000). The predicted number of clusters were tested using the silhouette R package to examine whether the clustering performed correctly. This approach resulted in 2 clusters for DAEs and epigenome features at 8-12 PCW, 3 clusters for 13-18 PCW and 2 clusters for >18 PCW, for each of CP and GZ, respectively. For each cluster, we determined the gene expression of protein coding genes interacting with the DAEs from each cluster, as obtained from published RNA-seq data sets. Significant differences in expression levels between different clusters were determined using the Wilcoxon signed rank test in R. Also, target genes linked to each cluster were used for functional enrichment analysis using Enrichr [109], as described under gene ontology analysis.

### Experimental validation

#### Cell culture

HEK293 LTV cells were cultured in DMEM medium (Gibco), supplemented with 10% FBS at 37°C, 5% CO2. Human neural stem cells (NSCs) (Gibco, a kind gift from Raymond Poot, Rotterdam) were cultured in NSC medium (KnockOut DMEM-F12 (Gibco), 2 mM L-glutamine (Gibco), 20 ng/ml bFGF (Peprotech), 20 ng/ml EGF (Peprotech), 2% StemPro Neural supplement (Gibco), 100U/ml penicillin and 100µg/ml streptomycin), as previously described [111].

#### Enhancer activity in STARR-seq plasmid

For experimental validation in **Figure 1G**, we selected 22 DAEs that showed interaction with a target gene by HiC. DAEs were amplified from genomic DNA and cloned into the STARR-seq plasmid (kind gift of A.Stark) [42] as previously described [51]. For the additional tested enhancer deletions (**Supplementary Figure 4**), the obtained STARR-seq plasmids containing *IRF2BPL*, *CHD2* and *MACF1* enhancers were modified by site-directed mutagenesis to remove regions with high or low ncER score. The following regions were deleted: *IRF2BPL* (chr14: 77422484-77422514); *CHD2* (ncER1 chr15: 93363603-93363640, ncER3 chr15: 93363780-93363790); *MACF1* (ncER1 chr1: 39598824-39598844, ncER2 chr1:39598744-39598754). The regions with low ncER score at the 5’ and 3’ ends (80-100bp) of *IRF2BPL*, *CHD2* and *MACF1* enhancers were excluded by Gibson assembly. Primer sequences are available upon request. HEK293 and NSC were transfected with STARR-seq plasmid containing enhancer regions using polyethylenimine (PEI, Sigma) or Lipofectamine™ Stem Transfection Reagent (Thermo Scientific) respectively. Spike-in of a pmCherry-N1 plasmid (Clonetech) was used as a transfection control. 24h post transfection cells were collected, stained with Hoechst dye and the enhancer activity was measured by FACS analysis (20,000 cells per sample). GFP-positive cells within the mCherry-positive population were quantified to assess enhancer activity compared to an empty STARR-seq vector. Two independent transfection experiments were performed, each in duplicates. Statistical analysis was performed using a one-way ANOVA test followed by multiple comparison test (Fisher’s LSD test). Calculations were conducted in GraphPad Prism v.8.

#### dCas9-KRAB-MeCP2 silencing of active enhancers in NSC

We selected DAEs linked to *CHD2*, *CAD* and *TRAK1* and designed for each DAE two targeting gRNAs (primer sequences are available upon request). gRNAs were cloned into a pgRGFP plasmid (Addgene #82695, a kind gift of Allan Mullen) [112]. NSCs were co-transfected with dCas9-KRAB-MeCP2 (Addgene #110824, kind gift of Alejandro Chavez and George Church) [113] and the two gRNAs/DAE and collected for RNA isolation 48h post transfection. Transfection efficiency was estimated by FACS analysis (78-92% GFP-positive cells detected). RNA was isolated using TRI reagent (Sigma) followed by cDNA preparation using iSCRIPT cDNA synthesis kit (BioRad). Fold change in gene expression (ΔΔct method) was evaluated by qPCR (iTaq universal SYBR Green Supermix) (Sigma), performed in CFX96RTS thermal cycler (Bio-Rad), as previously described [111].TBP expression was used as housekeeping normalization control. Statistical analysis was performed using a one-way ANOVA test followed by multiple comparison test (Fisher’s LSD test). Calculations were conducted in GraphPad Prism v.8.

#### Zebrafish studies

Selected DAEs used in the *in vitro* experiments were transferred by Gibson assembly between the AscI and PacI site of a E1b-GFP-Tol2 enhancer assay plasmid (a kind gift of Ramon Birnbaum) [114] containing an E1b minimal promoter followed by GFP, using the following transfer primers: Transfer_fw: 5’-AGATGGGCCCTCGGGTAGAGCATGCACCGG-3’ and Transfer_rv: 5’-TCGAGAGATCTTAATGGCCGAATTCGTCGA-3’. Constructs were injected into zebrafish embryos using standard procedures, together with Tol2 mRNA to facilitate genomic integration. At least 50 embryos were injected per construct in at least two different injection experiments. GFP expression was observed and annotated at 1, 2 and 3dpf by a fluorescent Leica M165FC stereomicroscope. Images were analyzed using imageJ (FIJI). An enhancer was considered active when at least 30% of the larvae showed consistent GFP expression.

### Data visualization

To generate UCSC Genome Browser Tracks, aligned reads were converted to bedgraph using genomeCoverageBed, after which the bedGraphToBigWig tool from the UCSC Genome Browser was used to create a bigwig file. All enhancer regions, Enhancer-Gene interactions and TAD coordinates were uploaded directly as bed files. Other plots were drawn using R packages.

## Declarations

### Ethics approval and consent to participate

For zebrafish studies, only larvae prior to 5 days post fertilization were used. Zebrafish were kept according to guidelines of the EMC animal welfare office

### Consent for publication

not applicable

### Availability of data and materials

All data generated in this study is supported by the included main and supplementary figures. Further supplementary details will be fully made available upon peer review. Some of the primary data that were used to support the findings of this study are available from dbGaP and PsychENCODE, but restrictions apply to the availability of these data, which were used under license for the current study and so are not publicly available (third party data). The related pipeline and processed data will be available after peer review or upon request.

### Competing interests

The authors declare that they have no competing interests.

### Funding

RD is supported by a China Scholarship Council (CSC) PhD Fellowship (201906300026) for her PhD studies at the Erasmus Medical Center, Rotterdam, the Netherlands. EM is supported by Netherlands Organisation for Scientific Research (ZonMW Off Road grant). TSB is supported by the Netherlands Organisation for Scientific Research (ZonMW Veni, grant 91617021), a NARSAD Young Investigator Grant from the Brain & Behavior Research Foundation, an Erasmus MC Fellowship 2017, and Erasmus MC Human Disease Model Award 2018. Funding bodies did not have any influence on study design, results and data interpretation or final manuscript.

### Authors’ contribution

SY performed primary computational analysis, with help of RD. KL, AN, and EP performed validation experiments in cells. EMS, HCvsL and TvH performed zebrafish validation experiments. SY, EM and TSB wrote the manuscript with input from all authors. EM and TSB conceived and jointly supervised the work.

## Acknowledgements

Ramon Birnbaum (Ben-Gurion University of the Negev, Israel) is acknowledged for sharing zebrafish reporter plasmids.

We would like to thank the following third parties for providing approved access to the indicated data sets that were used to support the findings presented in this study:

-phs000755.v2.p1: “BrainSpan Atlas of the Human Brain”. The datasets used for the analysis described in this manuscript were obtained from dbGaP at http://www.ncbi.nlm.nih.gov/gap through dbGaP accession number phs000406.v2.p1. Submission of the data, phs000406.v2.p1, to dbGaP was provided by Dr. Nenad Sestan. Collection of the data and analysis was supported by grants from the National Institutes of Health (MH089929, MH081896, and MH090047). Additional support was provided by the Kavli Foundation, a James S. McDonnell Foundation Scholar Award, NARSAD, and the Foster-Davis Foundation.

-phs000791.v1.p1: “Roadmap Epigenomics Program - UCSF”. Funding support for the NIH Roadmap Epigenomics Program was provided through the NIH Common Fund (Office of Strategic Coordination). Support for collection of datasets and samples was provided by a series of UO1 cooperative agreements with The Broad Institute [1U01ES017155-01], The Ludwig Institute for Cancer Research [1U01ES017166-01], The University of California San Francisco [1U01ES017154-01], and The University of Washington [1U01ES017156-01]. Data analysis and coordination were supported by an agreement with Baylor College of Medicine [1U01DA025956-01]. Assistance with data curation was supplied by GEO, and data access and visualization was supported by the NCBI.

-phs001226.v1.p1: “Regulatory Genomics of Human Embryonic Development”. The datasets used for the analysis described in this manuscript were obtained from dbGaP at http://www.ncbi.nlm.nih.gov/gap through dbGaP accession number phs001226.v1.p1.

-phs001438.v1.p1: “The dynamic landscape of open chromatin during human cortical neurogenesis”. Datasets from this study used for the analyses described in this manuscript were generated in the Geschwind laboratory and supported by NIH grants to D.H.G. (5R01MH060233; 5R01MH100027; 3U01MH103339; 1R01MH110927; 1R01MH094714). Datasets were obtained from dbGaP found at http://www.ncbi.nlm.nih.gov/gap through dbGaP study accession numbers phs001438.v1.p1.

-Data access from PsychENCODE was obtained for the following data sets: SynapseID: syn12033248 [9]; SynapseID: syn17092080 [34]. Data were generated as part of the PsychENCODE Consortium supported by: U01MH103339, U01MH103365, U01MH103392, U01MH103340, U01MH103346, R01MH105472, R01MH094714, R01MH105898, R21MH102791, R21MH105881, R21MH103877, and P50MH106934 awarded to: Schahram Akbarian (Icahn School of Medicine at Mount Sinai), Gregory Crawford (Duke), Stella Dracheva (Icahn School of Medicine at Mount Sinai), Peggy Farnham (USC), Mark Gerstein (Yale), Daniel Geschwind (UCLA), Thomas M. Hyde (LIBD), Andrew Jaffe (LIBD), James A. Knowles (USC), Chunyu Liu (UIC), Dalila Pinto (Icahn School of Medicine at Mount Sinai), Nenad Sestan (Yale), Pamela Sklar (Icahn School of Medicine at Mount Sinai), Matthew State (UCSF), Patrick Sullivan (UNC), Flora Vaccarino (Yale), Sherman Weissman (Yale), Kevin White (UChicago) and Peter Zandi (JHU).

---

**Supplementary Figure 1:**
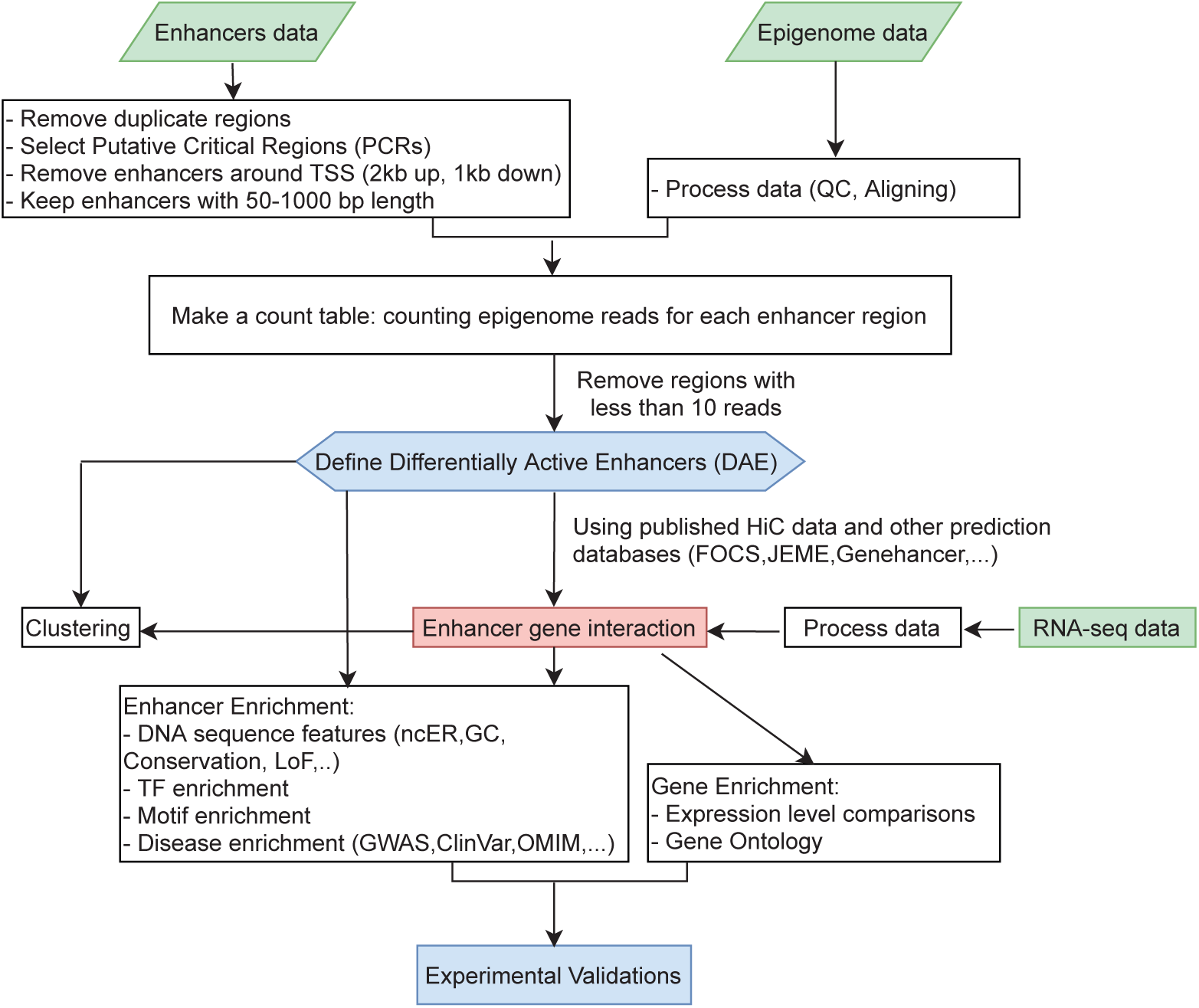
Flow chart of integrative data analysis. Overview of the various analysis steps performed in this study. See text and methods for additional details.

**Supplementary Figure 2:**
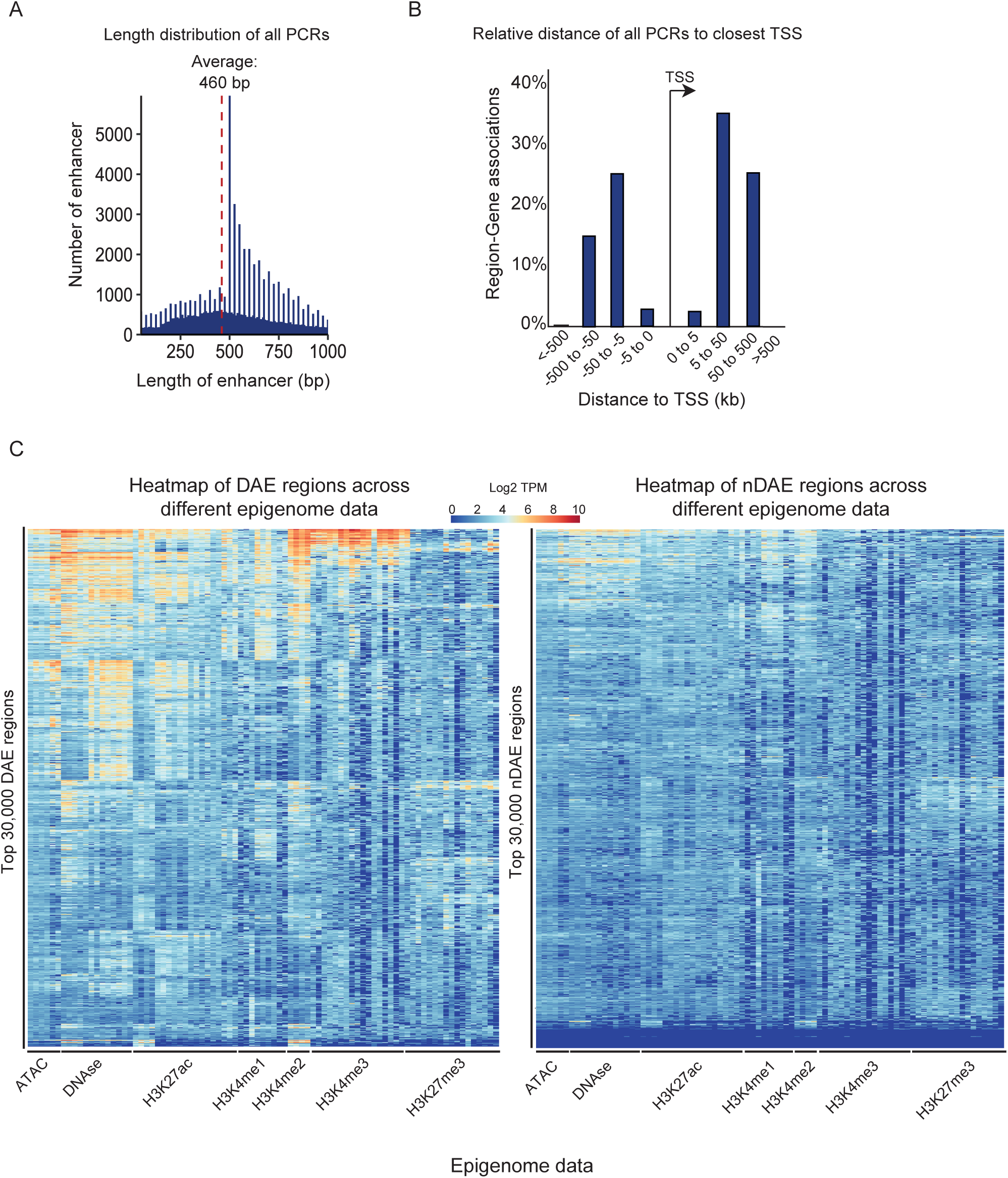
Derivation of pCRs and DAEs. A) Density plot showing the size distribution of the 202,163 pCRs in bps. The red dashed line indicates the average length of all pCRs (460 bp). B) Relative distribution of all 202,163 pCRs in relation to their closest transcriptional start site. Graph generated using GREAT [36]. C) Heatmaps showing variability across all epigenome features for the top 30,000 DAEs (left) and nDAEs (right). Columns represent in total 494 epigenome data sets used for the various types of histone marks and chromatin accessibility, as indicated.

**Supplementary Figure 3:**
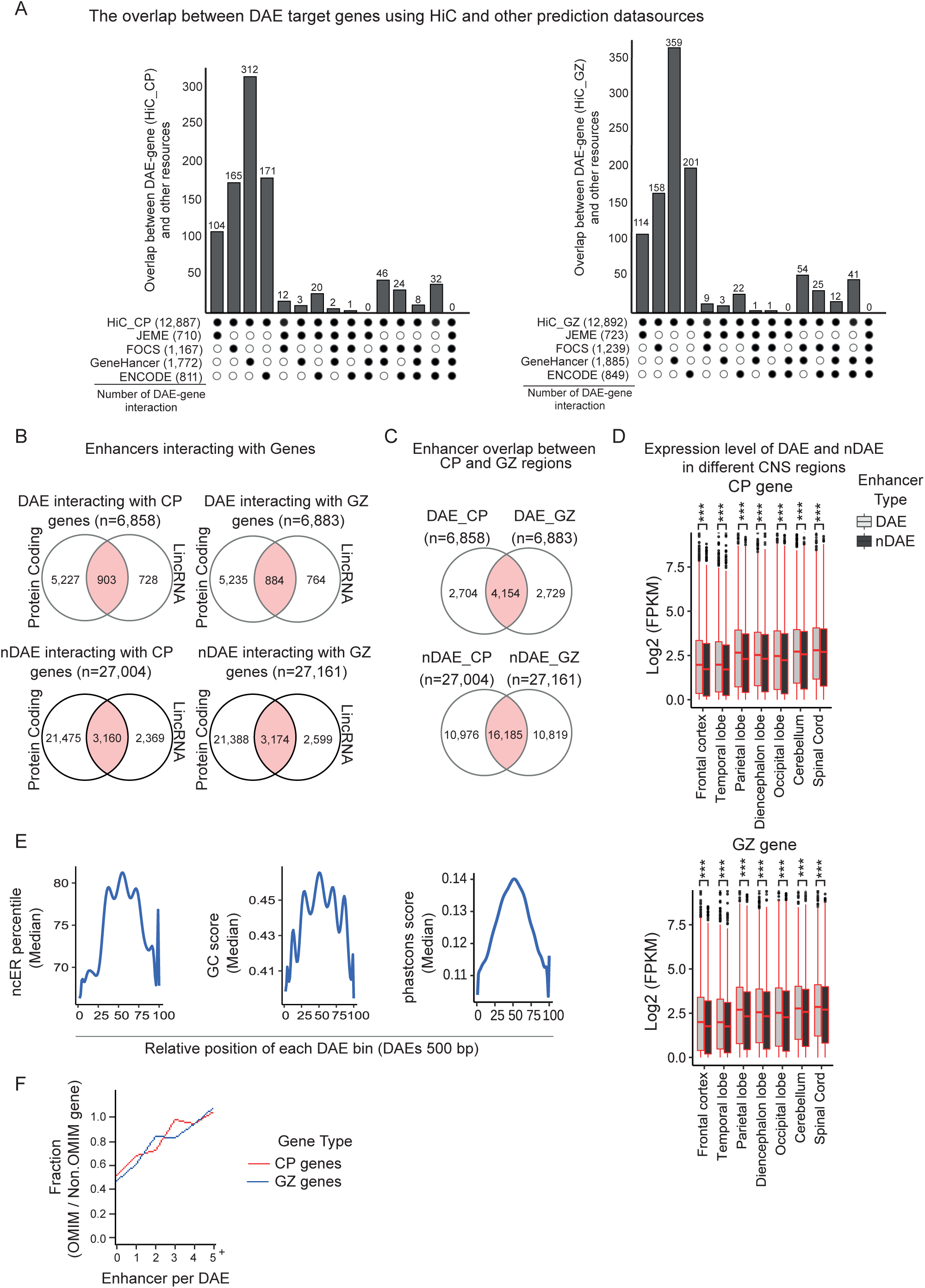
A) Bar chart showing the overlap between predicted enhancer-gene interactions from HiC of CP (left) or GZ (right) and the different other used enhancer-prediction methods JEME [37], ENCODE [38], FOCS [39] and GeneHancer [40]. B) Venn diagrams showing the interactions of DAEs (upper panel) or nDAEs (lower panel) with protein coding genes and lincRNA within the same TAD, for interactions from HiC in CP (left) or GZ (right). C) Venn diagrams showing the overlap between DAEs (upper panel) or nDAEs (lower panel) that interact with genes in CP (left) or GZ (right). D) Box plots showing the RNA-seq gene expression levels (in log2 FPKM) of genes linked to DAEs or nDAEs in CP (left) or GZ (right) for different brain regions. Boxes are interquartile range (IQR); line is median; and whiskers extend to 1.5 the IQR. *** p<0.001; (wilcox.test). RNA-seq data obtained from ENCODE project [7]. E) Line plot showing the distribution of the median ncER percentile (left) [43], GC content score (middle) [44] and phastcons score (right) [44] over all DAEs that have a size of 500 bp (n= 768). F) Line graph showing the fraction between OMIM divided by nonOMIM genes as a function of the number of enhancers that a DAE is interacting with, for interactions in CP (red) and GZ (blue). The more enhancers a DAE is interacting with, the more likely it is that the target gene of that DAE is a OMIM gene.

**Supplementary Figure 4:**
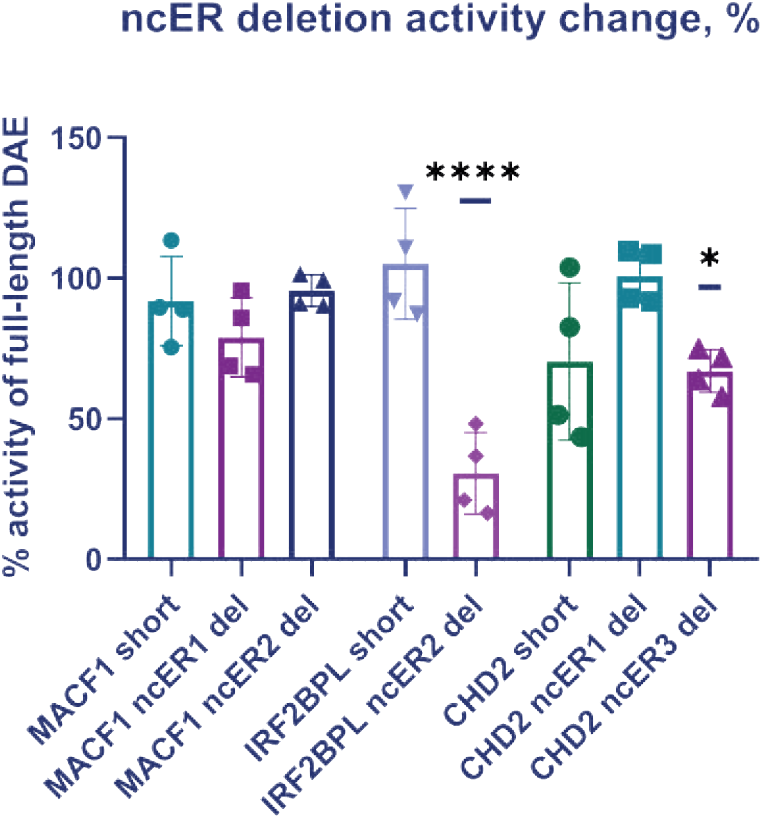
Effect of ncER deletion on activity of DAEs linked to IRF2BPL, CHD2 and MACF1. Percentage of activity of modified DAEs (see methods) compared to the full-length DAE in STARR-seq enhancer reporter experiments is plotted. Two independent transfection experiments were performed, each in duplicate. All data points and standard deviation are shown. * p<0.05; **** p<0.0001 (one-way ANOVA test followed by multiple comparison test (Fisher’s LSD test).

**Supplementary Figure 5:**
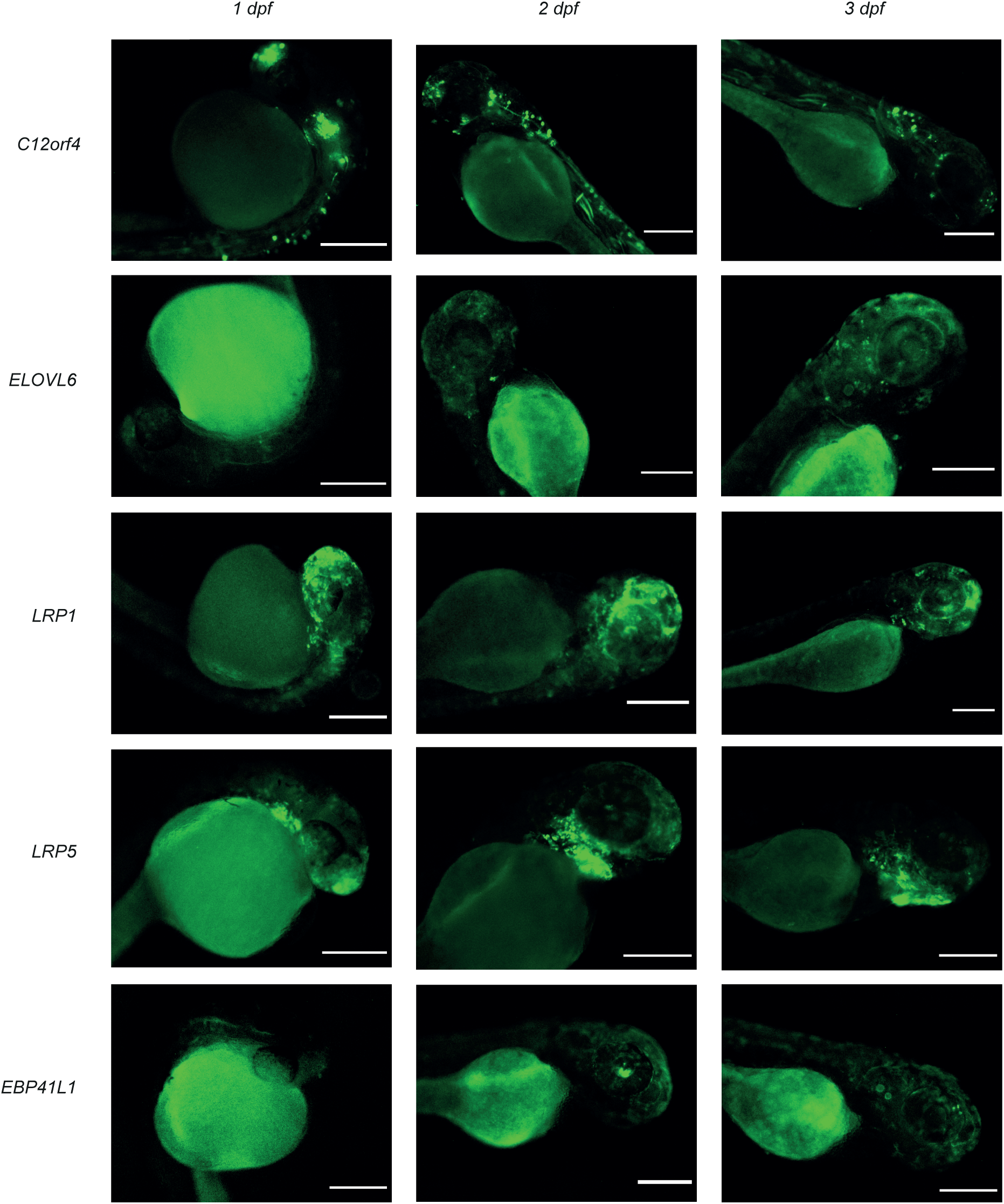
Zebrafish enhancer reporter assay. Panel of additional fluorescent images for validated enhancers, showing GFP expression in zebrafish at 1, 2 and 3 dpf. Scale bars represent 500 µm.

## Notes

### Competing Interest Statement

The authors have declared no competing interest.

